# Evaluation of genomic and phenomic prediction for application in apple breeding

**DOI:** 10.1101/2024.09.17.613481

**Authors:** Michaela Jung, Marius Hodel, Andrea Knauf, Daniela Kupper, Markus Neuditschko, Simone Bühlmann-Schütz, Bruno Studer, Andrea Patocchi, Giovanni AL Broggini

## Abstract

Apple breeding schemes can be improved by using genomic prediction models to forecast the performance of breeding material. These models’ predictive ability depends on factors like trait genetic architecture, training set size, relatedness of the selected material to the training set, and the validation method used. Alternative genotyping methods such as RADseq and complementary data from near-infrared spectroscopy could help improve the cost-effectiveness of genomic selection. However, the impact of these factors and alternative approaches on predictive ability beyond experimental populations still need to be investigated. In this study, we evaluated 137 prediction scenarios varying the described factors and alternative approaches, offering recommendations for implementing genomic selection in apple breeding. Our results show that extending the training set with germplasm related to the predicted breeding material can improve average predictive ability across eleven studied traits by up to 0.08. The study emphasizes the usefulness of leave-one-family-out validation, reflecting the application of genomic selection to a new family, although it reduced average predictive ability across traits by up to 0.24 compared to cross-validation. Similar average predictive abilities across traits indicate that imputed RADseq data could be a suitable genotyping alternative to SNP array datasets. The best-performing scenario using near-infrared spectroscopy data for phenomic prediction showed a 0.35 decrease in average predictive ability across traits compared to conventional genomic prediction, suggesting that the tested phenomic selection approach is impractical. These findings offer valuable guidance for applying genomic selection in apple breeding, ultimately leading to the development of breeding material with improved quality.

## INTRODUCTION

Genomic selection utilizes phenotyped and genotyped individuals to train genomic prediction models, enabling performance predictions for genotyped breeding material without available phenotypic information (Meuwissen et al., 2001). This approach has demonstrated the ability to increase genetic gain in breeding programs for major annual crops such as maize and wheat (Gaffney et al., 2015; Gaynor et al., 2017). For apple, an outcrossing perennial fruit crop, several genomic prediction studies have reported low to high predictive abilities, mostly dependent on the genetic architecture of traits and the population design (Kumar et al., 2012; Muranty et al., 2015; Migicovsky et al., 2016). Similarly, for the apple reference population, hereafter referred to as the apple REFPOP, the predictive ability is strongly dependent on the genetic architecture of the trait (Jung et al., 2020; Cazenave et al., 2021; Jung et al., 2022). The apple REFPOP was established in replicates in six European countries as a diverse population of apple accessions and European breeding material to facilitate implementation of genomic selection in local apple breeding programs (Jung et al., 2020). Phenotyping of the apple REFPOP resulted in the most extensive dataset of trait-environment combinations in apple to date, which has been instrumental for evaluating the accuracy of genomic prediction, dissecting the genetic architecture of traits, and identifying numerous marker-trait associations (Jung et al., 2022). These findings resulted in the recommendation for the adoption of genomic selection across the majority of the 30 examined quantitative traits (Jung et al., 2022). Despite extensive testing of genomic selection methodologies in apple, their applications have been scarce, with limited reports detailing practical implementation (Isik et al., 2015; Kostick et al., 2023).

The critical aspects for the practical integration of genomic selection in plant breeding lie in ensuring a substantial size of the training population and its close relatedness to the breeding material that is to be selected (Voss-Fels et al., 2019). As modest to strong improvements in predictive ability for apple traits have been shown by expanding the training population size (Cazenave et al., 2021; Minamikawa et al., 2024), enrichment of a training population composed of the apple REFPOP with local breeding material may positively influence predictive performance and allow accurate assessment of predictive ability for progenies closely related to future selection candidates.

To assess predictive ability, the widely adopted cross-validation has been applied in many studies evaluating genomic selection in apple (Kumar et al., 2012; Migicovsky et al., 2016; Jung et al., 2022). This method involves training models on a random subset of individuals and then estimating predictive ability for the remaining individuals in the validation set. Unlike cross-validation, the leave-one-family-out validation (LOFO) can verify the reliability of genomic prediction within a practical breeding scheme where phenotypic data for a specific family, aimed for genomic selection, is typically absent. Among the applications of LOFO in apple (Kumar et al., 2015; Minamikawa et al., 2021; Kostick et al., 2023), a study applying both LOFO and cross-validation showed generally lower values and higher variance of predictive ability for individual families used as validation sets in LOFO compared to cross-validation (Kostick et al., 2023). The assessment of the extent of decrease in predictive ability for LOFO compared to cross-validation remains to be assessed for the apple REFPOP and local breeding material.

When introducing genomic selection into a breeding program, an important consideration relates to the choice of the appropriate genotyping method. Single nucleotide polymorphism (SNP) array genotyping technologies allow repeated testing of germplasm for the same set of SNPs. Their applications in apple (Bianco et al., 2014, 2016) resulted in the acquisition of large-scale datasets for genomic prediction such as the apple REFPOP (Jung et al., 2020). Alternative genotyping technologies such as genotyping-by-sequencing (Elshire et al., 2011) or restriction site-associated DNA sequencing (RADseq) (Miller et al., 2007) capture new variation in each set of analyzed material. Genotyping-by-sequencing has proved efficient for genotyping and genomic prediction of apple germplasm collections (Migicovsky et al., 2016, 2022). Even in the presence of considerable amounts of missing data, the alternative genotyping technologies are recognized for their cost-effectiveness and sufficient informativeness, and they could present a viable alternative for genotyping apple breeding material. However, the novel SNP variation and missing data may pose a challenge when integrating datasets from alternative genotyping technologies with those from SNP arrays, and the potential of such a combination should be evaluated.

In the search for cost-efficient predictive solutions, the concept of phenomic prediction based on near-infrared spectroscopy (NIRS) data could provide an alternative to genomic prediction (Rincent et al., 2018). Building prediction models using NIRS data alone or combined with genomic data have demonstrated predictive abilities comparable to or even surpassing those of genomic data alone for various quantitative traits in annual crops such as wheat and soybean (Rincent et al., 2018; Zhu et al., 2021). However, phenomic prediction have often shown lower predictive abilities than genomic prediction for perennial species such as grapevine and poplar (Rincent et al., 2018; Brault et al., 2022). The potential of phenomic prediction for accurate predictions of quantitative apple traits remains to be estimated.

Aiming for practical and cost-effective prediction of apple traits in biparental families, this study evaluated the predictive ability of eleven quantitative traits using genomic prediction models based on (i) apple REFPOP and apple REFPOP enriched with the Swiss breeding material (denoted as AZZ material, with the abbreviation AZZ derived from the project name) from three breeding programs (Agroscope, Lubera, Poma Culta) for model training, (ii) LOFO and cross-validation for model validation using apple REFPOP families and AZZ families, and (iii) SNP arrays and an alternative genotyping technology based on RADseq for genotyping. Finally, (iv) the aim was also to compare the concepts of phenomic prediction with genomic prediction (including their combination) in assessing the predictive ability of the eleven quantitative traits within the accessions of the apple REFPOP.

## MATERIALS AND METHODS

### Plant material

The apple REFPOP contained 265 progenies from 27 biparental families (∼10 genotypes per family) produced by several European breeding programs and 270 diverse accessions (Jung et al., 2020). Five locations of the apple REFPOP were considered for this study, namely (i) Rillaar, Belgium, (ii) Angers, France, (iii) Laimburg, Italy, (iv) Lleida, Spain, and (v) Waedenswil, Switzerland. All genotypes were replicated at least twice at every location and planted in 2016 following a randomized complete block design. Three control genotypes ‘Gala’, ‘Golden Delicious’, and ‘CIVG198’ were replicated up to 22 times at each location. The genotypes were grown under the agricultural practice common to each location (integrated plant protection).

The AZZ material was composed of 390 progenies from 23 biparental families (17 genotypes per family on average). Additionally, the AZZ material included 234 advanced selections from the Swiss breeding programs of Agroscope (141), Lubera (53) and Poma Culta (45), and five accessions important as founders for the studied breeding programs. All genotypes of Agroscope were assumed to share their location with the apple REFPOP due to proximity of the orchards in Waedenswil, Zurich, Switzerland. Progenies of Lubera were grown in Buchs, St. Gallen, Switzerland. The advanced selections and accessions of Lubera were located in Felben, Thurgau, Switzerland. No control genotypes were present at the locations of the Lubera breeding program. All genotypes of Poma Culta, including individual trees of the control genotypes ‘Gala’, ‘Golden Delicious’ and ‘Rustica’/’Rusticana’, the latter control genotype also present among the accessions at Agroscope, were grown in Hessigkofen, Solothurn, Switzerland. All AZZ progenies were part of the early stages of the three Swiss breeding programs and therefore not clonally replicated. The AZZ advanced selections and accessions were replicated 2–18 times per location depending on the stage of selection and plant material availability. Tree replicates were grown side by side. All trees were planted between 2010 and 2020. The trees grown at Agroscope and Lubera were managed according to integrated plant protection management, while the trees at Poma Culta were grown according to biodynamic plant protection management.

### Phenotyping

Eleven traits (floral emergence, harvest date, flowering intensity, total fruit weight, number of fruits, single fruit weight, titratable acidity, soluble solids content, fruit firmness, red over color, and russet frequency) were scored in the apple REFPOP for up to five years during 2018–2022. For the AZZ material, the phenotyping of the eleven traits was done during two years in 2021– 2022 and, when available, historical data were retrieved for 2018–2020. In the apple REFPOP, all traits were estimated from individual trees, i.e., genotype replicates. To assess titratable acidity, soluble solids content, fruit firmness, and russet frequency, a random sample of 5–20 fruits was drawn for each tree. For the AZZ material, the traits floral emergence, harvest date, flowering intensity, total fruit weight, number of fruits and single fruit weight were scored for individual trees, and the traits titratable acidity, soluble solids content, fruit firmness, red over color, and russet frequency were measured from pooled samples of 5–20 fruits across the available trees of each genotype.

Floral emergence was estimated in Julian days as the date when the first 10% of flowers opened. Flowering intensity was scored on a nine-grade scale as the percentage of existing flowers from the maximum possible number of flowers. Fruits were harvested on harvest date, when at least 50% of the fruit on a tree was fully mature, which was determined in Julian days based on fruit ripening estimated by expert knowledge. Total fruit weight per tree was measured in kilograms (kg) and all fruits were counted to assess the number of fruits. Single fruit weight in grams (g) was calculated as the ratio of the total fruit weight to the number of fruits. Titratable acidity (g/kg), soluble solids content (°Brix) and fruit firmness (g/cm^2^) were measured within one week after the harvest date using an automated laboratory Pimprenelle (Setop, France). Red over color was the percentage of red fruit skin assessed on a 6-grade scale. Russet frequency was the proportion of fruits with russet skin in the fruit sample. Total fruit weight, number of fruits, single fruit weight, red over color, and russet frequency were evaluated at harvest. Additional details about the assessment of the eleven traits can be found in Jung et al. (2022).

### Genotyping using SNP arrays

The apple REFPOP was genotyped for biallelic SNPs using a combination of the Illumina Infinium® 20K SNP genotyping array (Bianco et al., 2014) and the Affymetrix Axiom® Apple 480K SNP genotyping array (Bianco et al., 2016) as described by Jung et al. (2020). From the 270 apple REFPOP accessions and 265 progenies, 269 accessions and 6 progenies were genotyped using the 480K SNP array. These data as well as additional genomic data obtained with the 480K SNP array for 1,089 genotypes representing a diversity collection of accessions were retrieved from previous studies (Urrestarazu et al., 2017; Muranty et al., 2020) to be used as reference set for genotype imputation, and they were available at a resolution of 303,239 SNPs (Jung et al., 2020). For the remaining 259 apple REFPOP progenies, 20K SNP array genomic data of 7,054 SNPs were obtained from previous studies (Bianco et al., 2014; Bink et al., 2014; Laurens et al., 2018; Jung et al., 2020). The 20K SNP array was additionally used to genotype 390 progenies and 239 advanced selections and accessions of the AZZ material as well as the apple REFPOP accession and control genotype ‘CIVG198’. The resulting dataset of 6,863 SNPs was derived by filtering according to the iGLMap (Di Pierro et al., 2016) and retaining SNPs whose physical positions matched those of 303,239 SNPs from the 480K SNP array. All physical SNP positions were based on the doubled haploid GDDH13 (v1.1) reference genome (Daccord et al., 2017), and all SNPs from the 20K SNP array were also found in the 480K SNP array set. Missing genotype information in the 20K SNP array data was imputed to reach the resolution of 303,239 SNPs as described by Jung et al. (2020). For the imputation, pedigree information (Howard et al., 2018; Muranty et al., 2020) together with the imputation set of genotypes comprised of all available 480K SNP array data were supplied to the software Beagle (v4.0) (Browning and Browning, 2007). Finally, the SNP array dataset contained 303,239 biallelic SNPs for 1,164 progenies, advanced selections, and accessions of the apple REFPOP and AZZ material.

### Restriction site-associated DNA sequencing

DNA was extracted from freeze-dried leaf tissue using a custom DNA extraction kit (LGC, Germany) for 181 AZZ progenies (samples). An adapted double digest RADseq (ddRADseq) protocol from Peterson et al. (2012) was applied using the EcoRI-HF and TaqI-v2 restriction enzymes (New England Biolabs, MA, USA). Illumina NovaSeq 6000 paired-end sequencing was used to produce reads of 150 bp length. Additional details about the applied RADseq genotyping approach can be found in Online Resource 1.

Raw sequences were demultiplexed and reads with any uncalled base were removed using Stacks (v2.59) (Catchen et al., 2013). Adapter sequences were trimmed and reads shorter than 36 bp were removed using HTStream (v1.3.0) (https://github.com/s4hts/HTStream). The reads were aligned to the doubled haploid GDDH13 (v1.1) reference genome (Daccord et al., 2017) using Bowtie2 (v2.4.4) (Langmead and Salzberg, 2012). Variants were called using BCFtools (v1.15.1) (Danecek et al., 2021) applying the commands “mpileup” and “call” with a minimum base quality of 20. The following variant filtering was inspired by Migicovsky et al. (2022). VCFtools (v0.1.16) (Danecek et al., 2011) was used to retain genotypes with a minimum depth of 4 reads (per sample and variant) and variants with the mean depth of at least 4 reads (per variant over all samples). The software PLINK (v1.9-beta6.18) (Chang et al., 2015; Purcell and Chang, n.d.) was used to remove (i) indels and multiallelic variants, (ii) SNPs with minor allele frequency lower than 0.01, (iii) SNPs unassigned to the 17 apple chromosomes, (iv) SNPs with missing call rates exceeding 0.7 and (v) samples with missing call rates exceeding 0.7. The variant filtering required removal of 12 samples with poor genotyping quality. One sample was additionally removed due to a possible mistake at sampling. The RADseq dataset finally consisted of 281,558 SNPs for 168 samples (AZZ progenies).

Out of the 281,558 SNPs of the RADseq dataset, 7,255 SNPs overlapped in their physical positions with the 303,239 SNPs of the SNP array dataset. Using these 7,255 overlapping SNPs, the missing genotypic information in the RADseq dataset was imputed. Prior to the imputation, reference allele mismatches in the genotypes of the SNP array dataset were determined and the strand orientation was corrected using the plugin “fixref” of BCFtools (v1.15.1) (Danecek et al., 2021). BCFtools was further used to remove two SNPs with duplicated physical position from the SNP array dataset, decreasing its size from 303,239 to 303,237 SNPs. The imputation was performed as described by Jung et al. (2020) using the software Beagle (v4.0) (Browning and Browning, 2007) with pedigree information (Howard et al., 2018; Muranty et al., 2020) and 1,364 genotypes of the SNP array dataset that were originally genotyped applying the 480K SNP array. The imputed RADseq dataset consisted of 303,237 SNPs for 168 samples (AZZ progenies).

The 168 genotypes present in both the imputed RADseq dataset and the SNP array dataset were compared to evaluate the imputation performance using the command “stats” of BCFtools (v1.15.1) (Danecek et al., 2021). Non-reference discordance rate was defined for each genotype as the ratio of mismatches among the three possible allele dosages (0, 1, 2) to the sum of these mismatches, heterozygous matches, and homozygous alternative matches. The squared Pearson correlation between the allele dosages of the compared datasets was calculated for each genotype. Non-reference discordance rate and the squared Pearson correlation were averaged over all 168 genotypes.

### Near-infrared spectroscopy

To obtain the NIRS dataset for the apple REFPOP accessions, 20 fresh leaves were collected from one replicate of each of the 263 accessions. These samples were taken between August 15 and 23, 2022, in the integrated plant protection part of the apple REFPOP orchard in Waedenswil, Switzerland. The leaves were collected from sun-exposed branches located at least 0.5 m above ground, dried at 60°C for approx. 48 h, then stored with silica gel until milling using a centrifugal mill (Retsch, Germany). NIRS measurements of wavenumbers 10,500–4,000 cm^-1^ (wavelength 950–2500 nm) were taken per milled leaf sample (i.e., tree) at three different locations on the leaf powder-filled measurement cell using the NIRFlex N-500 spectrometer (Büchi, Switzerland). The three NIRS measurements were averaged per tree, then the wavelengths of 950–1049 nm were removed due to inconsistencies to produce a matrix of the raw NIRS. The normalized NIRS were obtained from the raw NIRS by scaling and centering to the mean of 0 and standard deviation of 1. Detrended NIRS were estimated from the raw NIRS applying a standard normal variate transformation followed by fitting a second order linear model using the R package prospectr (v0.2.6) (Barnes et al., 1989; Stevens and Ramirez-Lopez, 2022). Derivatives of the raw and normalized NIRS were estimated applying a Savitzky-Golay smoothing filter using the R package signal (v0.7-7) (https://gnu-octave.github.io/packages/signal/). For the first derivative, a filter order of 2 and filter length of 37 were applied. For the second derivative, the filter order of 3 and filter length of 61 were used. Finally, seven matrices for raw and pre-processed NIRS with wavelengths of 1050–2500 nm represented by 1,380 variables in each matrix were obtained for further analyses.

### Population structure analysis

Principal component analysis was performed for 1,164 genotypes of the SNP array dataset using the R package FactoMineR (v2.4) (Lê et al., 2008). For the same dataset, population network was visualized using the so-called NetView approach as described in detail by Neuditschko et al. (2012) and Steinig et al. (2016). Briefly, genetic distances were computed by subtracting pairwise identical-by-state (IBS) relationships, as provided by PLINK (v1.9) (Chang et al., 2015; Purcell and Chang, n.d.), from 1. This algorithm was applied using the number of k nearest neighbors k-NN = 30. To illustrate the genetic relatedness between neighboring genotypes, the thickness of the edges (connecting lines) was associated with the proportion of genetic distance, with thicker edges corresponding to smaller genetic distances. To identify highly informative genotypes in the population network, the genetic contribution score (gc_j_) was calculated for each genotype and the node size was scaled accordingly. The computation of the gc_j_ was based on Singular Value Decomposition (SVD) of a symmetric relationship matrix and accounted for the correlation between the j-th individual relationship vector and the i-th standardized eigenvector, limiting the number of eigenvectors to the first k significant principal components (Neuditschko et al., 2017). Therefore, the aforementioned IBS relationships were converted into a symmetric relationship matrix, and the number of k significant principal components was determined with the modified version of Horn’s parallel analysis described by Glorfeld (1995) and implemented in the R package paran (v1.5.3). Here, a significance level of p = 0.01 was used and the number of iterations was set to 10,000.

### Phenotypic data analysis

Statistical data analysis was carried out with the available multi-environmental phenotypic data separately for each of the eleven traits. This was to ensure good quality of the phenotypic data, decompose variance into various effects and estimate clonal means as well as clonal mean heritability. For these analyses, the environment was defined as a combination of location and year. The raw phenotypic values for the apple REFPOP and AZZ material were used to estimate environment-specific clonal mean heritability (Jung et al., 2022). Trait-environment combinations with the environment-specific clonal mean heritability value of less than 0.1 were removed. The orchard design of the apple REFPOP allowed to correct the raw phenotypic values for spatial heterogeneity when modelling spatial trends using two-dimensional P-splines using the R package SpATS (v1.0-11) (Rodríguez-Álvarez et al., 2018). The spatial correction was done separately for every combination of trait and environment to produce adjusted phenotypic values of each tree as described by Jung et al. (2020). The adjusted phenotypic values of each tree were denoted here as the adjusted tree values. The adjusted tree values for the apple REFPOP and the raw phenotypic values for the AZZ material (spatial correction not possible due to design) were used to fit a linear mixed-effects model using the R package lme4 (v1.1-23) (Bates et al., 2015)

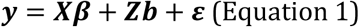

where 𝒚 was a vector of the phenotypic values, 𝑿 the design matrix for the fixed effects, 𝜷 the vector of fixed effects, 𝒁 the design matrix for the random effects, 𝒃 the vector of random effects and 𝜺 the vector of random errors. The model included a fixed effect of environment and the random effects of genotype and genotype by environment interaction. An additional fixed effect of tree age was used for traits floral emergence, harvest date, flowering intensity, total fruit weight, number of fruits and single fruit weight, where the phenotypic values were obtained for individual trees in both the apple REFPOP and AZZ material. The model fit was initially used for outlier detection following Bernal-Vasquez et al. (2016). After the identified outliers were removed, the model was refitted to extract the conditional means of the random effect of genotype (clonal values). Pearson correlations and their significance tests were computed between clonal values for all pairs of traits. The model fit was further used to obtain the phenotypic variance explained by the random effects of genotype 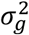 and the genotype by environment interaction 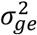 as well as the error variance 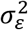. The estimated variances were used to assess the across-environmental clonal mean heritability as the genotypic variance 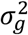 over the phenotypic variance 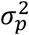. The phenotypic variance was calculated as

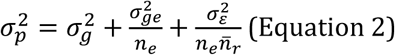

where 𝑛_𝑒_ was the number of environments and 𝑛̅_𝑟_ the mean number of genotype replications. Additionally, the fraction of phenotypic variance associated with the fixed effects was estimated as the variance of the vector predicted from the model fit when all random effects were set to zero. For 263 apple REFPOP accessions with NIRS data available, the clonal values were re-estimated for five sets of environments: (i) Waedenswil, 2020 (one environment), (ii) Waedenswil, 2021 (one environment), (iii) Waedenswil, 2022 (one environment), (iv) Waedenswil, 2018–2020 (five environments), (v) five apple REFPOP locations, 2018–2020 (all available environments). For subsets of adjusted tree values from one environment, the model following Equation 1 included only the random effect of genotype. For subsets of adjusted tree values from five or more environments, the model following Equation 1 included the fixed effect of environment and the random effects of genotype and genotype by environment interaction. In case of singular model fit, the random effect of genotype by environment interaction was dropped from the model.

### Genomic prediction

The model genomic-BLUP (G-BLUP) was applied to make genomic predictions for progenies from biparental families. It was based on a genomic relationship matrix 𝑮, which resulted from the cross-product of centered and standardized SNP values divided by the number of SNPs in the SNP matrix (VanRaden, 2008). The model was defined as

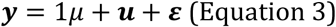

where 𝒚 was a response vector of the clonal values for one trait, 𝜇 was an intercept, 𝒖 was a vector of random effects following 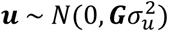 with variance 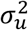, and 𝜺 was a vector of residuals assuming 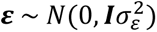. The G-BLUP was fitted using a semi-parametric Bayesian reproducing kernel Hilbert spaces regression algorithm implemented in the R package BGLR (v1.0.8) (Pérez and de los Campos, 2014). The G-BLUP was applied with 12,000 iterations of the Gibbs sampler, a thinning of five, and a burn-in of 2,000 discarded samples. Predictive ability was estimated as Pearson correlation coefficient between clonal values and predicted values for the validation set of progenies from biparental families.

#### Genomic prediction based on the SNP array dataset

The SNP array dataset of 303,239 SNPs was used to construct the genomic relationship matrix 𝑮 and fit the model G-BLUP following Equation 3. Different combinations of training and validation sets for LOFO were defined: (i) whole apple REFPOP but one family as the training set, the excluded apple REFPOP family as the validation set, the model was fitted separately for each of the 27 apple REFPOP families, (ii) whole apple REFPOP as the training set, 23 AZZ families in a single validation set (i.e., not a LOFO in a strict sense), (iii) whole apple REFPOP and AZZ material but one apple REFPOP family as the training set, the excluded apple REFPOP family as the validation set, the model was fitted separately for each of the 27 apple REFPOP families, and (iv) whole apple REFPOP and AZZ material but one AZZ family as the training set, the excluded AZZ family as the validation set, the model was fitted separately for each of the 23 AZZ families. For each of these four combinations, predictive ability was estimated (i) specifically for each validation family (LOFO1), and (ii) across all validation families (LOFO2), resulting in eight prediction scenarios.

Three additional prediction scenarios were implemented using a ten-fold cross-validation repeated ten times, resulting in 100 model runs for each scenario. The genotypes were split into folds randomly without replacement. The scenarios were defined as: (i) 90% of the apple REFPOP genotypes as the training set and 10% of the apple REFPOP genotypes as the validation set, with predictive ability estimated for the apple REFPOP families, (ii) 90% of the apple REFPOP and AZZ genotypes as the training set and 10% of the apple REFPOP and AZZ genotypes as the validation set, with predictive ability assessed for the apple REFPOP families, and (iii) same as (ii) but predictive ability assessed for the AZZ families. Predictive ability was estimated separately for each of the ten repetitions of the ten-fold cross validation (across ten folds), resulting in ten values of predictive ability per scenario and trait.

#### Genomic prediction based on the RADseq dataset

The model G-BLUP following Equation 3 was used to estimate the impact of genotyping method (RADseq or SNP array) on the predictive ability for progenies from biparental families. The SNP array dataset was used for model training, with the training set consisting of the entire apple REFPOP and AZZ material but one AZZ family (LOFO). Validation sets were constructed from 168 AZZ progenies across 13 families with RADseq data available. These progenies were divided into 13 distinct validation sets, each corresponding to a single family. Model validation was conducted using either the imputed RADseq dataset or the SNP array dataset, and the models were fitted with three different sets of SNPs: (i) full set of 303,237 SNPs, (ii) subset of 7,255 SNPs that overlapped in their physical positions between the RADseq dataset and the SNP array dataset, and (iii) subset of 7,255 randomly sampled SNPs from the entire set of 303,237 SNPs. For each of these six prediction scenarios, i.e., combinations of two validation data types (imputed RADseq or SNP array datasets) and three sets of SNPs, predictive ability was estimated across genotypes of all 13 validation families (LOFO2). This resulted in one estimation of predictive ability for each scenario and trait, except in the case of the third set of SNPs, where the random sampling was repeated 20 times, resulting in 20 estimations of predictive ability for each trait.

### Comparison of phenomic and genomic prediction

Genomic and phenomic prediction, as well as their combination, were compared to assess the potential advantages of integrating NIRS data for predicting apple traits across 263 apple REFPOP accessions. First, the genomic prediction model was implemented following Equation 3, with the genomic relationship matrix 𝑮 based on 303,239 SNPs of the SNP array dataset. Second, the phenomic prediction models were constructed following a similar framework to the genomic prediction, with the distinction that the 𝑮 matrix was substituted by one of the seven phenomic relationship matrices {𝑯_𝟏_, 𝑯_𝟐_, …, 𝑯_𝟕_}, which were derived from raw and pre-processed NIRS matrices. Third, the combined prediction models incorporated the matrix 𝑮 in combination with one of the phenomic relationship matrices. The combined prediction models followed the Equation 3 with the random effects defined as 𝒖 = 𝒖_𝟏_ + 𝒖_𝟐_ where 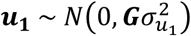 with variance 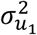 and 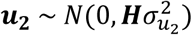 with variance 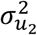. 𝑯 was a generic representation of one of the phenomic relationship matrices {𝑯_𝟏_, 𝑯_𝟐_, …, 𝑯_𝟕_}. In all three types of prediction models, the response vector 𝒚 represented either one of the re-estimated clonal values for five sets of environments or the adjusted tree values from 2020, 2021 or 2022, obtained for the same trees as those used for NIRS measurements. The combinations of prediction models (one genomic, seven phenomic and seven combined) with eight different response vectors 𝒚 resulted in 120 prediction scenarios.

For each prediction scenario and trait, a ten-fold cross-validation was repeated ten times, with 263 apple REFPOP accessions split into the folds randomly without replacement, resulting in 100 model runs. Every model run comprised 90% of the accessions as the training set and 10% of the accessions as the validation set. Predictive ability was estimated once across all apple REFPOP accessions in each ten-fold cross-validation, resulting in ten values of predictive ability for each prediction scenario and trait.

Unless otherwise specified, all statistical analyses and data formatting in this work were conducted using R (v3.6.0 and v4.2.2) (R Core Team, 2022), and the graphical visualizations were created using the R package ggplot2 (v3.4.0) (Wickham, 2016).

## RESULTS

### Characteristics of the genomic datasets

Combining the marker sets obtained with the 480K SNP array (303,239 SNPs for the apple REFPOP accessions and the diversity collection of accessions) and the 20K SNP array (7,054 SNPs for apple REFPOP progenies, and 6,863 SNPs for the AZZ advanced selections and accessions as well as AZZ progenies, both SNP sets imputed to the resolution of 303,239 SNPs) resulted in the SNP array dataset of 303,239 SNPs for 2,253 genotypes (Fig. 1). Acquired as an alternative to SNP arrays for a genotype subset, the raw RADseq output was characterized by the mean number of reads per sample of 3,313,087 (minimum 2,051, maximum 9,636,664), the mean alignment rate of 86.12% (minimum 74.71%, maximum 89.64%), and 1,394,979 raw variants. Filtering of these variants resulted in the RADseq dataset consisting of 281,558 SNPs for 168 AZZ progenies (Fig. 1). The density of markers along the chromosomes in the RADseq and SNP array datasets was similar (Fig. S1 in Online Resource 2). The physical positions of 7,255 SNPs overlapped between the RADseq and SNP array datasets (Fig. 1b), and the distribution of overlapping SNPs along the chromosomes was less consistent compared to the entire RADseq and SNP array datasets (Fig. S1 in Online Resource 2). Based on the overlapping SNPs, the imputation of the RADseq dataset at the physical positions matching the SNP array dataset resulted in the imputed RADseq dataset of 303,237 SNPs. A comparison of 168 AZZ progenies found in both the imputed RADseq dataset and the SNP array dataset showed the average non-reference discordance rate of 27.82 and the average squared Pearson correlation between the allele dosages of the compared datasets of 0.74.

**Fig. 1.**
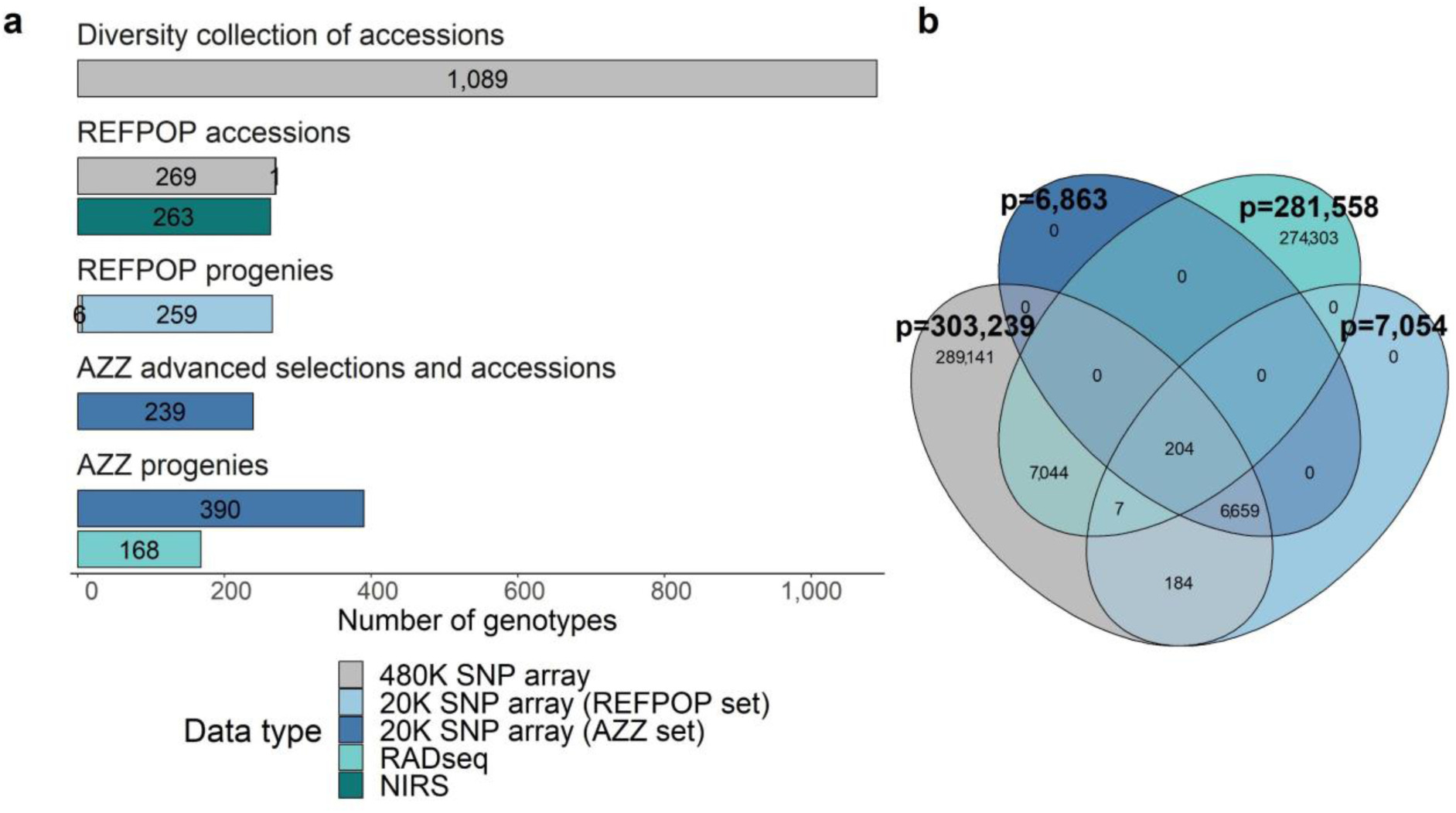
Overview of the datasets. **a** Three genotyping approaches (480K SNP array, 20K SNP array, RADseq) and near infrared spectroscopy (NIRS) were used to score various types of genotypes (accessions, progenies, advanced selections) of the diversity collection, apple REFPOP (REFPOP), and AZZ material (AZZ). **b** Venn diagram illustrating the extent of overlap in physical SNP positions between the RADseq dataset, and three sets of SNPs utilized in generating the SNP array dataset (non-bold numbers). Numbers in bold indicate the total number of SNPs (p) in each set. Colors in b correspond to legend in a (excluding NIRS).

### Characteristics of the phenotypic datasets

Of all the genotyped material, 1,164 progenies, advanced selections, and accessions of the apple REFPOP and AZZ material were phenotyped for eleven traits. Groups of material based on its origin (apple REFPOP or AZZ material) and location showed similarities in their distributions between phenotyping years (Fig. 2a, Fig. S2 in Online Resource 2). Among the studied traits, the largest proportion of the phenotypic variance explained by the genotypic effects of 63.93% and 65.70% was found for harvest date and red over color, respectively (Fig. 2b). The same traits also exhibited the highest values of across-environmental clonal mean heritability of 0.99 and 0.98 for harvest date and red over color, respectively (Fig. S3 in Online Resource 2). Flowering intensity showed the lowest proportion of the phenotypic variance explained by the genotypic effects of 4.69% and the lowest across-environmental clonal mean heritability of 0.71 (Fig. 2b, Fig. S3 in Online resource 2). High positive significant correlation of 0.82 was observed between total fruit weight and number of fruits (Fig. S4 in Online Resource 2).

**Fig. 2.**
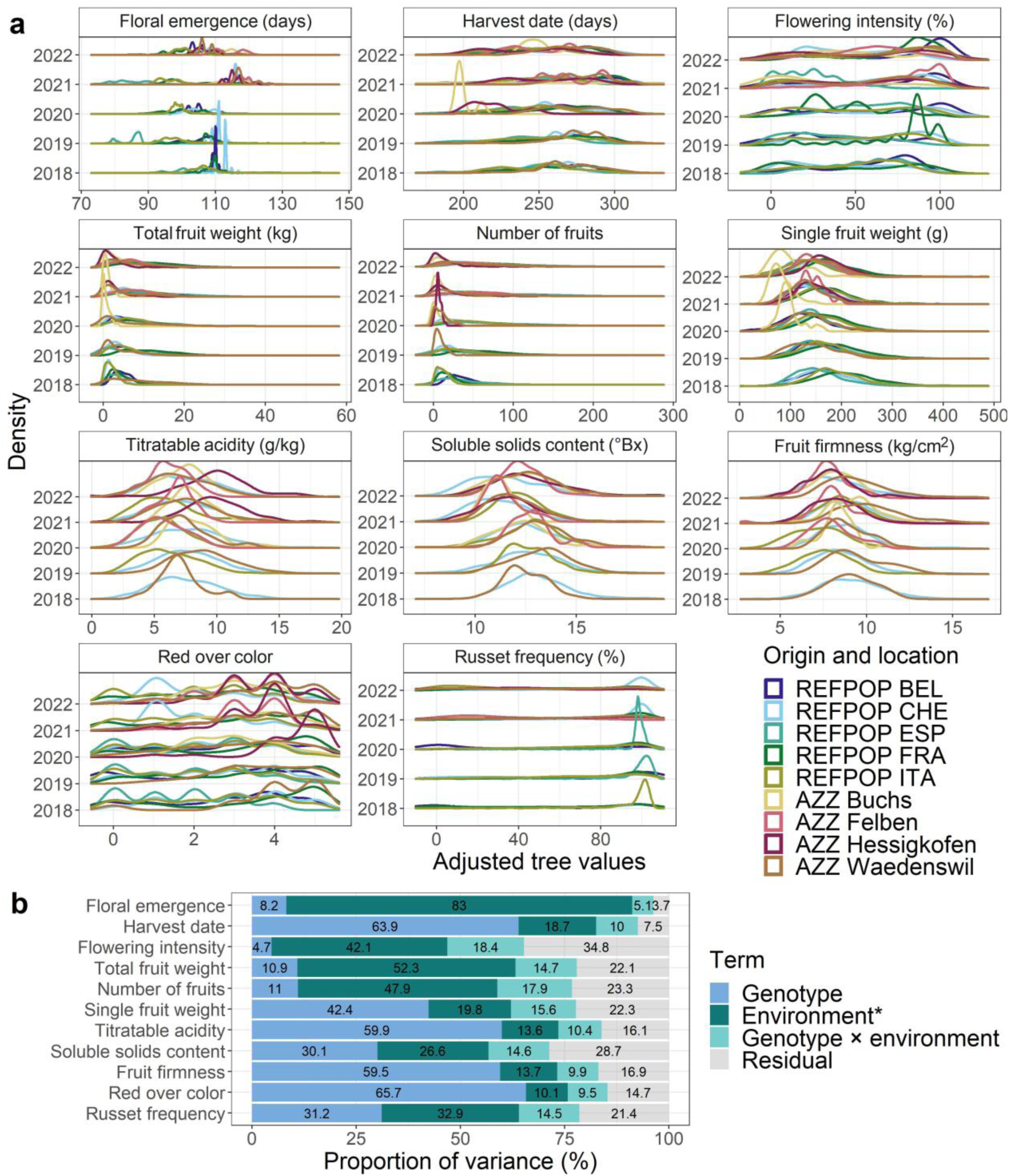
Characteristics of the phenotypic dataset composed of eleven quantitative traits. **a** Distributions of traits shown as density estimates separately for each year of phenotyping, colored by a combination of origin and location of material. The abbreviations stand for: apple REFPOP (REFPOP), AZZ material (AZZ), Belgium (BEL), Switzerland (CHE), Spain (ESP), France (FRA) and Italy (ITA). **b** Proportion of variance explained by the effects of genotype, environment*, genotype × environment interaction and residual. *The proportion of the effect of environment was based on fixed effects of environments (titratable acidity, soluble solids content, fruit firmness, red over color, and russet frequency) or the fixed effects of environments and tree age (floral emergence, harvest date, flowering intensity, total fruit weight, number of fruits, and single fruit weight).

### Population structure analysis

Principal component analysis (Fig. 3a) showed that the first two principal components explained 7.34% of the total variance in the SNP array dataset. The first two principal components illustrated a strong overlap among groups of material from various sources, including apple REFPOP and AZZ material from the breeding programs of Agroscope, Lubera, and Poma Culta. An exception was observed among the apple REFPOP accessions, which were partially positioned apart from the remaining material along the first principal component. Moreover, AZZ progenies from Lubera formed a separate cluster distinguished along the second principal component. These weaker relationships between the apple REFPOP accessions and the remaining material, as well as between the AZZ progenies from Lubera and the remaining material, were even more evident in the population network (Fig. 3b). Additionally, groups of AZZ progenies from Agroscope and Poma Culta were separated from the material allocated in the center of the population network, identifying additional population substructures. The node sizes of the population network, associated with gc_j_ (derived based on 37 significant principal components accounting for 87.77% of the genetic variance) highlighted ‘Gala’, ‘Golden Delicious’, and ‘Topaz’ as the accessions accounting for the greatest proportion of variance among all the studied material.

**Fig. 3.**
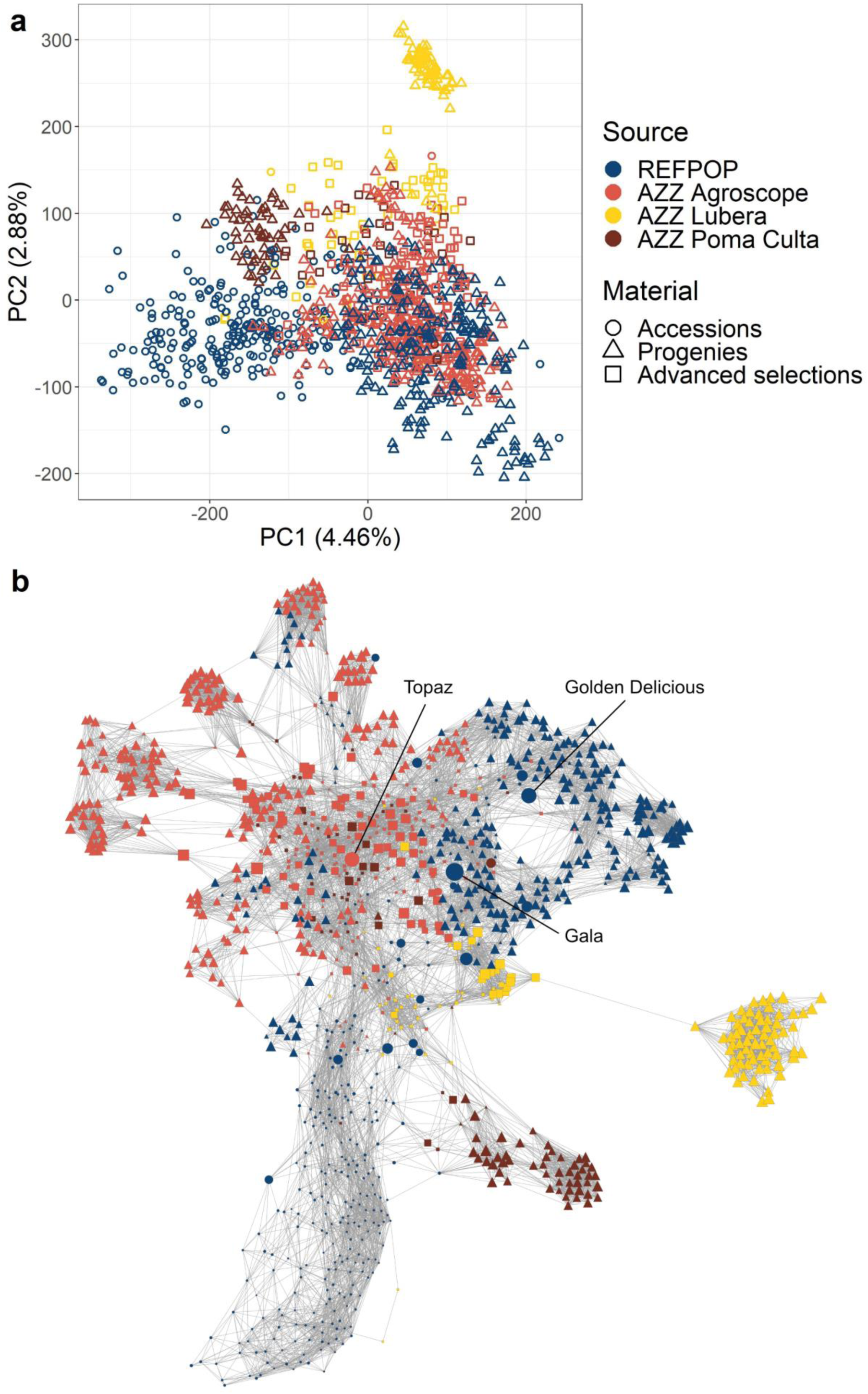
Population structure analysis for the SNP array dataset with 1,164 accessions, progenies and advanced selections of the apple REFPOP and AZZ material from the breeding programs of Agroscope, Lubera and Poma Culta. **a** First two principal components from the principal component analysis. **b** Population network, where each accession is illustrated by a node, with individual node size proportional to genetic contribution score (gcj). The thickness of edges varies in proportion to genetic distances, enabling the visualization of individual relationships within the dataset. Colors and shapes correspond to legend in a.

### Genomic prediction of biparental families

The genomic prediction of biparental families, using the SNP array dataset and applying LOFO or cross validation, structured into eleven prediction scenarios, resulted in predictive abilities that were highly variable across the studied traits (Fig. 4, Table S1 in Online Resource 3). Combining apple REFPOP with AZZ material for model training resulted in a higher average predictive ability across traits compared to using training sets consisting of apple REFPOP alone (Fig. 4, Table S1 in Online Resource 3), the increase in average predictive ability across traits for validation sets of apple REFPOP progenies and AZZ progenies were as follows: 0.05 and 0.04 in LOFO1, and 0.03 and 0.08 in LOFO2, respectively. Using cross-validation, this comparison of training sets resulted in an increase of 0.02 in average predictive ability across traits for validation sets of apple REFPOP progenies. In the case of validation sets of AZZ progenies, predictive abilities were not estimated using cross-validation for training sets consisting only of apple REFPOP, as AZZ progenies were not available for predictive ability estimation in this scenario.

**Fig. 4.**
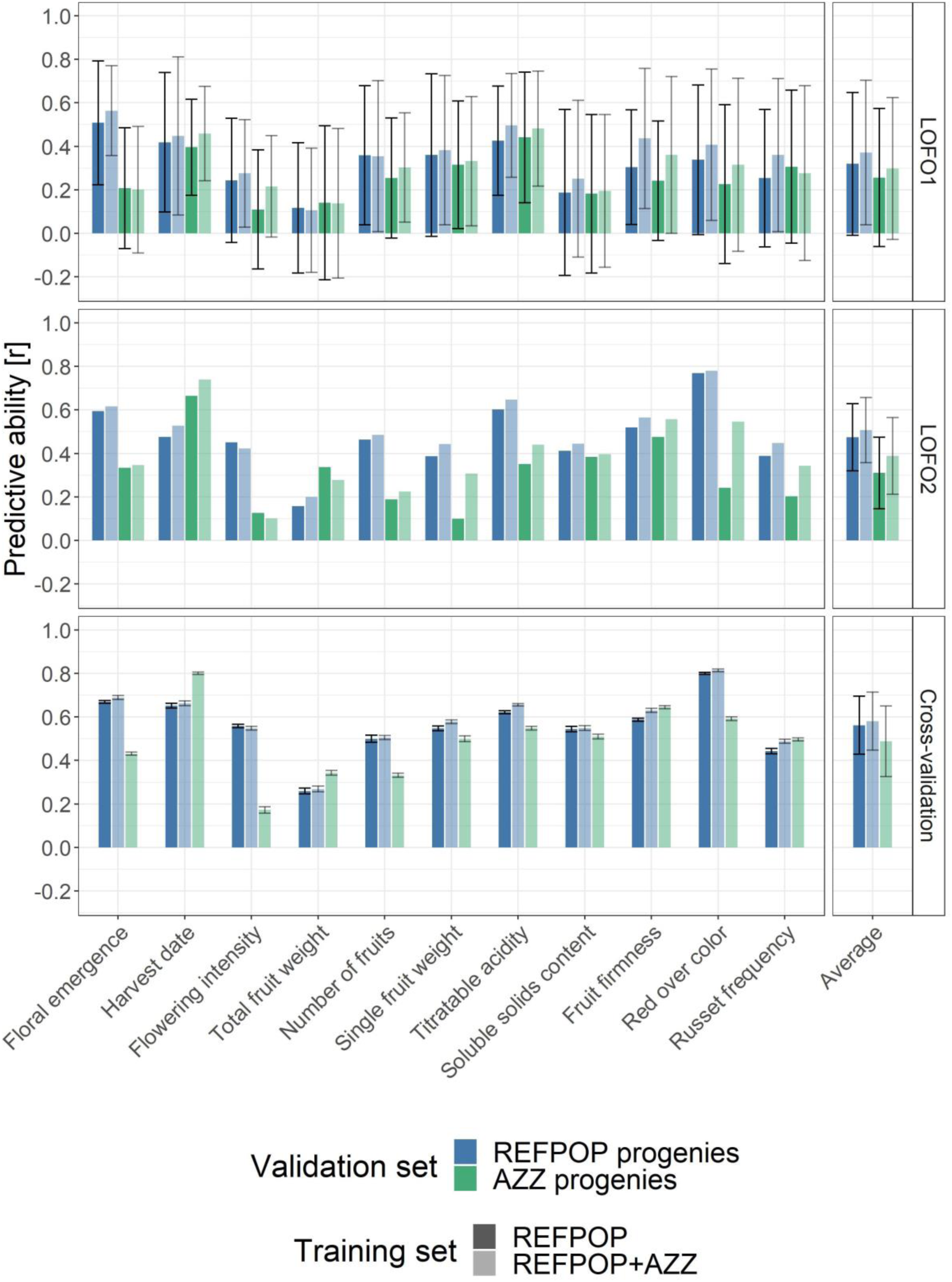
Genomic prediction of biparental families based on the SNP array dataset of 303,239 SNPs. Combinations of two training sets (opacity) and two validation sets (color) were used to assess predictive ability (PA) for eleven studied traits. Leave-one-family-out model validation was used to estimate predictive ability per family (LOFO1, top) or across families (LOFO2, center). Cross-validation was deployed to assess predictive ability across families (bottom). AZZ progenies were not available for predictive ability estimation using cross-validation based on training sets consisting only of apple REFPOP. Error bars correspond to standard deviation around the mean.

Validation sets composed of apple REFPOP progenies showed higher average predictive ability across traits than validation sets comprising AZZ progenies (Fig. 4, Table S1 in Online Resource 3). Applying LOFO1, the difference in average predictive ability across traits between the validation sets composed of apple REFPOP progenies and AZZ progenies was 0.06 for training sets composed solely of apple REFPOP and 0.07 for training sets composed of both apple REFPOP and AZZ material. In LOFO2, this difference increased to 0.12 and 0.16, respectively. The difference was 0.09 using cross-validation for training sets composed of apple REFPOP combined with AZZ material.

When predictive ability was estimated specifically for each validation family using LOFO1, high variability of predictive abilities was observed within each trait and on average over traits (Fig. 4, Table S1 in Online Resource 3). The average predictive abilities across traits estimated over all validation families using LOFO2 were higher than for LOFO1, with the range of the average predictive ability increase varying from 0.05 to 0.15 between scenarios. An additional increase in average predictive ability across traits was observed for the prediction scenarios implementing cross-validation with predictive abilities across families, the increase ranging from 0.07 to 0.10. Genomic prediction of validation families based on imputed RADseq dataset as an alternative to the SNP array dataset resulted in average predictive abilities for individual traits and across traits that showed strong differences between the compared prediction scenarios (Fig. 5, Table S2 in Online Resource 3). When imputed RADseq data were provided for the validation family and the model was trained with the subset of 7,255 SNPs that overlapped in their physical positions between the RADseq dataset and the SNP array dataset, a near-zero average predictive ability across traits of 0.01 was obtained. This predictive ability increased to 0.15 for the identical subset of 7,255 SNPs when, instead of the imputed RADseq data, the SNP array data were provided for the validation family. The imputed RADseq data for the validation family resulted in an improved average predictive ability across traits of 0.31 when the entire SNP set of 303,237 SNPs was utilized for model training. For the full SNP set of 303,237 SNPs, the average predictive ability across traits increased to 0.34 when the SNP array dataset was used for model validation. Additional two prediction scenarios comparing SNP sets of 7,255 SNPs that were randomly sampled among the 303,237 SNPs of the SNP array dataset resulted in similar predictive abilities between the validation data types and compared to the full set of 303,237 SNPs (Table S2 in Online Resource 3).

**Fig. 5.**
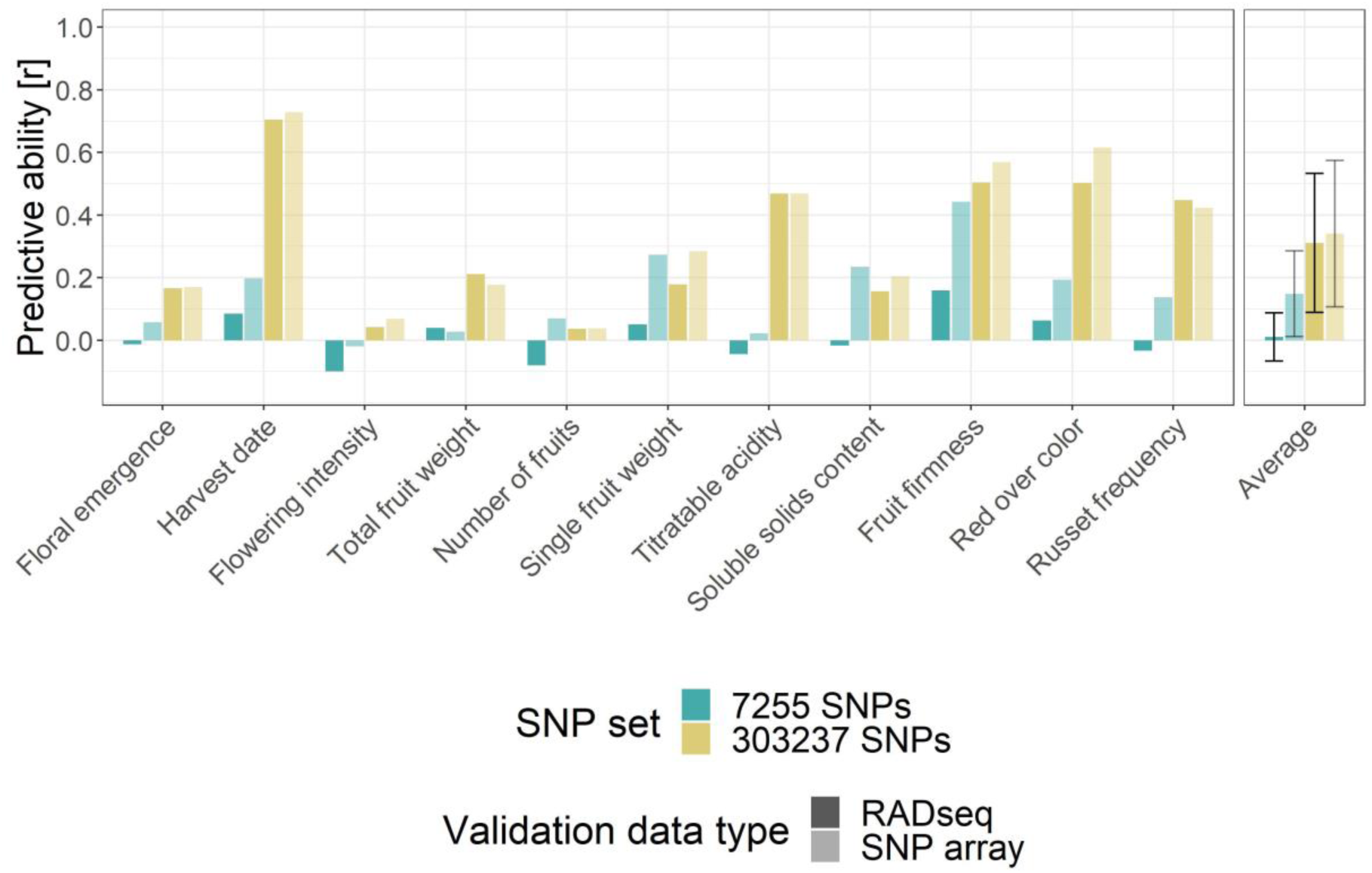
Genomic prediction of biparental families based on the imputed RADseq dataset as an alternative to the SNP array dataset. Genomic prediction models were trained for eleven traits using the SNP array dataset for the entire apple REFPOP and AZZ material but one validation family. Model training was repeated using SNP sets of different extent (color), with 7,255 SNPs representing the overlap in physical SNP positions between the RADseq dataset and SNP array dataset, and 303,237 SNPs being the full extent of the imputed RADseq and SNP array datasets. For the validation family, the imputed RADseq or SNP array data were provided (opacity). Predictive ability for each trait was estimated across all validation families (LOFO2). For the averaged predictive abilities across traits, the error bars correspond to standard deviation around the mean.

### Phenomic and genomic prediction of accessions

Using cross-validation, the evaluation of 120 prediction scenarios representing combinations of prediction models (seven phenomic, one genomic, and seven combined) with eight different response vectors showed that the best performing scenario reached an average predictive ability across traits of 0.56 (Fig. 6, Fig. S5 in Online Resource 2, Table S3 in Online Resource 3). This scenario was based on genomic prediction using the clonal values estimated across all available environments (clonal values, 5 locations, 2018-2022). Following closely behind this scenario was the best performing combined model integrating raw NIRS alongside SNPs together with the clonal values estimated across all available environments, exhibiting a decrease in average predictive ability across traits of 0.003 compared to the overall best model. Among the scenarios implementing models for phenomic prediction, the highest average predictive ability across traits of 0.21 was found for the design matrix consisting of the first derivative of normalized NIRS and the response vector comprising the adjusted tree values from the site and year of the NIRS measurement (Waedenswil, 2022).

**Fig. 6.**
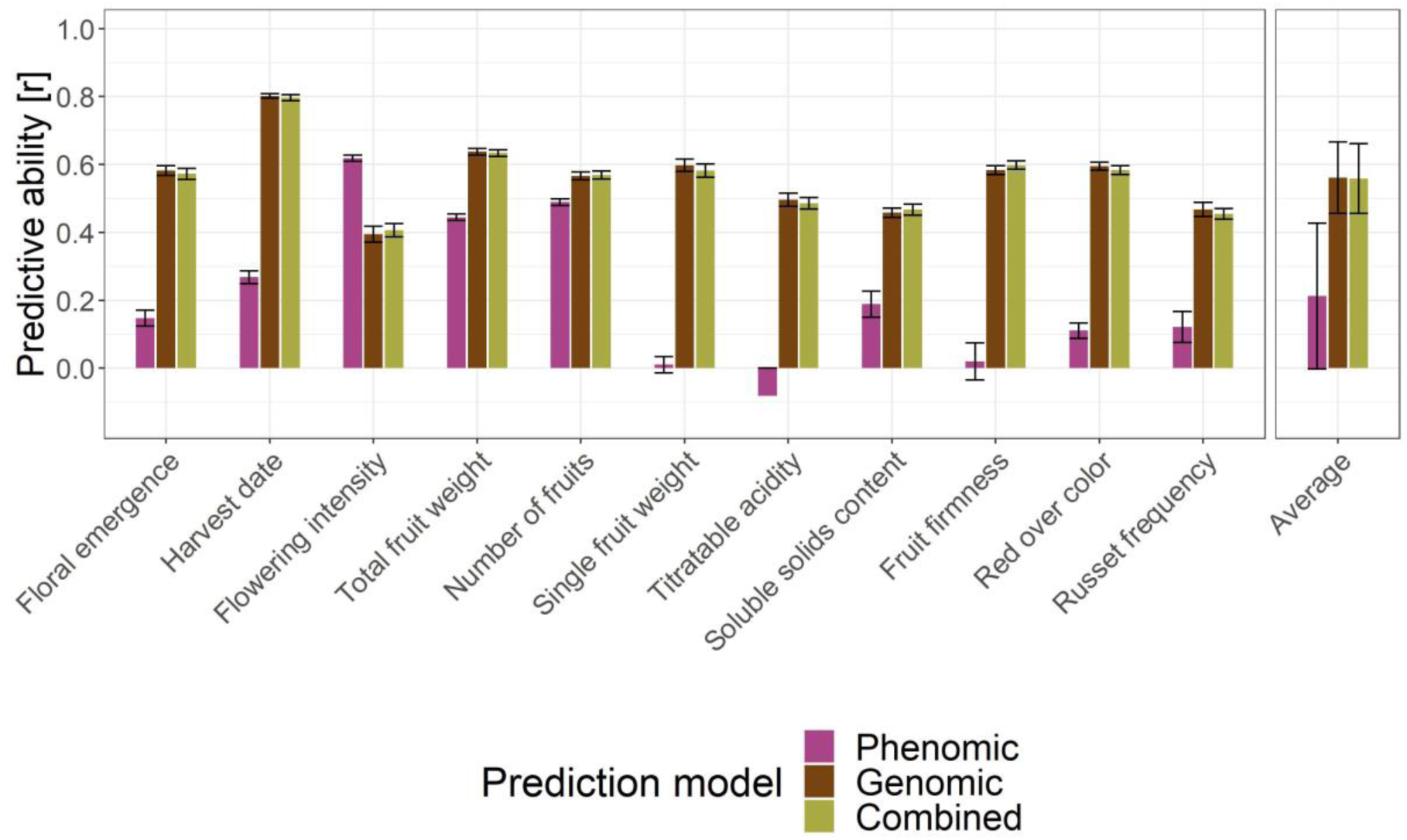
Predictive ability of the phenomic prediction, genomic prediction, and their combination, assessed by cross validation for eleven traits using the apple REFPOP accessions. For each prediction model type (color), the visualized best performing prediction scenarios were: (i) phenomic prediction based on the first derivative of normalized NIRS and the response vector of the adjusted tree values from the site and year of the NIRS measurement (Waedenswil, 2022), (ii) genomic prediction based on the SNP array dataset and the response vector of the clonal values for all available environments (up to five locations and five years), and (iii) the combined prediction using a combination of raw NIRS with the SNP array dataset and the same response vector as in scenario ii. Error bars correspond to standard deviation around the mean.

The comparison of the best performing scenarios for phenomic, genomic, and combined prediction models revealed that the genomic and combined prediction models outperformed the phenomic prediction model for most traits (Fig. 6). Only for flowering intensity, the phenomic prediction model exhibited a higher predictive ability of 0.62 compared to the genomic and combined prediction models, which showed average predictive abilities of 0.39 and 0.41, respectively. In addition to flowering intensity, the phenomic prediction model achieved moderately high predictive abilities for other productivity traits (total fruit weight and number of fruits). However, phenomic prediction did not surpass genomic and combined prediction for these traits. The combined prediction model showed only minimal or no improvement in average predictive ability for individual traits compared to the genomic prediction model.

## DISCUSSION

This study addresses practical aspects of genomic selection application, including the composition of training and validation sets, differences between validation approaches, and the exploration of RADseq as an alternative genotyping technology for the prediction of quantitative traits in biparental families. It also presents the first comparison in apple between phenomic prediction, genomic prediction, and their combined approach for predicting performance of apple accessions.

### Impact of training set enlargement on population structure and predictive ability

To increase the size of the training population and ensure close relatedness to future breeding material that is targeted for genomic selection, the apple REFPOP dataset was extended with Swiss breeding material from the programs of Agroscope, Lubera, and Poma Culta (AZZ material). Population structure analysis showed that much of the apple REFPOP breeding material (progenies) was closely related to the progenies and advanced selections of the AZZ material (Fig. 3). This strong relatedness translated into a low to moderate increase in predictive abilities when training sets combined apple REFPOP with AZZ material, compared to training sets composed solely of apple REFPOP (Fig. 4). The increase in predictive ability was more pronounced for validation sets of AZZ progenies, with an average predictive ability across traits increasing by up to 0.08, likely due to increased relatedness between the training and validation sets achieved by incorporating AZZ material into the training set.

Despite recommendations for very large training populations that are closely related with the selection candidates (Voss-Fels et al., 2019), combining populations does not always increase predictive ability (Hayes et al., 2009; Cazenave et al., 2021). The modest increase in predictive ability observed when combining populations in our study may be attributed to the already accurate model training using the apple REFPOP alone, given the high relatedness found between genotypes of apple REFPOP and the predicted biparental families of the AZZ material.

Some genotypes included in the combined population of apple REFPOP and AZZ material showed weaker relatedness to the core of this population, as illustrated in the population network (Fig. 3b). Progenies from the breeding programs of Lubera and Poma Culta showed weaker relatedness, possibly due to the focus of these programs on specialized markets (home garden and biodynamic fruit farming, respectively). Additionally, a substantial part of the apple REFPOP accessions formed a distinct group at the bottom of the population network, with individual genotypes explaining only a small proportion of the genetic variance. This suggests that not all apple REFPOP accessions may be necessary for accurately predicting traits in the studied European and Swiss breeding material. Optimizing the training population by including fewer accessions could potentially provide similar predictive abilities while reducing phenotyping expenses (Akdemir and Isidro-Sánchez, 2019).

The largest proportion of explained variance in the population network for the accessions ‘Gala’, ‘Golden Delicious’, and ‘Topaz’ highlighted them as most important genotypes among the studied material (Fig. 3b). The traditional varieties ‘Gala’ and ‘Golden Delicious’ are widely cultivated (https://ec.europa.eu/eurostat/statistics-explained/index.php?title=Agricultural_production_-_orchards#Apple_trees) and closely related, with ‘Gala’ being a progeny of ‘Golden Delicious’. The modern variety ‘Topaz’, whose grandparent is ‘Golden Delicious’, has also gained popularity, not only for its production qualities, but also for its resistance to apple scab conferred by the resistance gene *Rvi6* (Gessler and Pertot, 2012). The success of these varieties in cultivation contributed to their repeated use in European and Swiss apple breeding.

### Differences between validation sets of progenies

Predictive ability was assessed separately for the validation sets of apple REFPOP progenies and AZZ progenies due to differences in their origin, experimental design, management, and years of phenotypic assessment. The apple REFPOP had five years of phenotyping conducted across five European locations applying a randomized complete block design, contributing to highly precise clonal values. In contrast, the AZZ material had an experimental design partially lacking randomization, replication, and control genotypes, used different management practices, and was phenotyped only in two consecutive years, probably resulting in lower quality clonal values. Additionally, some AZZ material, made for specialized markets, showed lower relatedness to the combined population of apple REFPOP and AZZ material (Fig. 3). These factors likely contributed to the lower average predictive ability across traits for AZZ progenies compared to apple REFPOP progenies, the differences in average predictive ability across traits ranging from 0.06 to 0.16 (Fig. 4).

Different trait variability between apple REFPOP progenies and AZZ progenies may also explain the differences in predictive ability. For example, red over color is a highly heritable trait that showed very high predictive ability in the apple REFPOP, where classes with lower proportions of red over color were well represented (Jung et al., 2022). Unlike apple REFPOP, the AZZ material predominantly consisted of genotypes with intensely red-colored skin (Fig. S2 in Online Resource 2). This decreased phenotypic variability in the AZZ material could contribute to weaker correlations between clonal values and predicted values, resulting in decreased predictive ability.

### Validation approach effect on predictive ability

When comparing LOFO1 (predictive ability estimated per validation family), LOFO2 (predictive ability estimated over all validation families), and the more commonly used cross-validation, the latter resulted in higher average predictive ability across traits, with increases of up to 0.10 compared to LOFO2 and up to 0.24 compared to LOFO1 (Fig. 4). However, it can be argued that the lower estimates of predictive ability from the LOFO approaches more accurately reflect the practical accuracy of genomic prediction. This is because cross-validation involves random splitting of validation families into training and test sets, which allows for model training using phenotypic information that would not be available in a practical apple breeding program.

A study focused on practical aspects of genomic selection in apple by Kostick et al. (2023) showed predictive abilities that were comparable to ours for prediction scenarios using LOFO1 and cross-validation, particularly for similarly measured traits such as single fruit weight, titratable acidity, soluble solids content, and red over color. In both studies, the average predictive ability across these traits was 0.54 when using cross-validation. This was observed in our study for the prediction scenario that included validation sets of AZZ progenies and training sets of apple REFPOP and AZZ material. When comparing this value to the average predictive ability across the four traits obtained using LOFO1, a decrease of 0.21 was found in our work and 0.13 in the study by Kostick et al. (2023). The lower average predictive ability for LOFO1 was associated with even more pronounced variability in predictive abilities for individual traits in our study compared to Kostick et al. (2023). A potential explanation for these differences in average predictive abilities between the compared studies lies in phenotypic variability. Since phenotypes of individuals from biparental families tend to cluster around the parental mean, there is low variability in clonal values and predictive abilities within the same family. This can reduce the ability of the correlation coefficient to detect systematic relationships between clonal and predicted values, especially in small families where random fluctuations may have a large impact. Therefore, the small average family size of 17 genotypes in the AZZ progenies likely contributed to less accurate and more variable predictive abilities for individual validation families using LOFO1. Indeed, family sizes of 80 to 100 genotypes were found to be more suitable for within-family prediction (Kostick et al., 2023). However, when all AZZ progenies were pooled for predictive ability estimation in LOFO2, the family sizes used allowed for similarly accurate genomic prediction as found by Kostick et al. (2023) using larger family sizes and LOFO1, with a decrease in average predictive ability across the four traits by 0.12 compared to cross-validation (Fig. 4). This comparison demonstrates the impact of phenotypic variability on estimating predictive ability. It also suggests that low estimates of predictive ability do not necessarily indicate inaccurate predictions.

### RADseq as an alternative genotyping technology to SNP arrays

Various amounts of SNPs have been generated from genotyping-by-sequencing pipelines for genomic prediction in apple, ranging from 6,400 to 98,584 SNPs (Migicovsky et al., 2016; McClure et al., 2018, 2019; Kumar et al., 2020). Our RADseq dataset, which included 281,558 SNPs for 168 AZZ progenies, was comparable in SNP count to the dataset of 278,224 SNPs obtained by genotyping-by-sequencing for 1,175 accessions in Canada’s Apple Biodiversity Collection (Migicovsky et al., 2022). Moreover, the overlap of 7,255 SNPs between our RADseq and SNP array datasets was comparable in SNP count to the previously reported overlap of 7,060 SNPs between the apple REFPOP datasets from 20K and 480K SNP arrays (Jung et al., 2020). The extent of this overlap allowed for the application of the imputation approach previously used for the apple REFPOP (Jung et al., 2020) to impute missing SNP information in the current RADseq dataset.

Genomic predictions of biparental families using LOFO2 performed with the imputed RADseq dataset of 303,237 SNPs showed similar average predictive abilities across traits as genomic predictions using 303,237 SNPs of the SNP array dataset, as well as 7,255 randomly sampled SNPs from the 303,237 markers of the SNP array dataset (Fig. 5, Table S2 in Online Resource 3). These results indicate that the applied imputation approach effectively filled in missing information in the RADseq dataset, enabling successful integration of the studied SNP datasets for accurate genomic prediction. However, our results also suggested that the uneven distribution along chromosomes of the 7,255 SNPs overlapping between the RADseq and SNP array datasets (Fig. S1 in Online Resource 2) hindered genomic predictions, as shown by the average genomic predictive ability across traits approaching zero when these SNPs alone were used for training genomic prediction models (Fig. 5). Replacing the imputed RADseq dataset by the SNP array dataset at the physical positions of the overlapping 7,255 SNPs, the average genomic predictive ability across traits was slightly restored to 0.15 (Fig. 5), pointing to genotyping errors in the RADseq dataset. Overall, our study demonstrated that genotype imputation using a reference set of high-density SNP array data is crucial for overcoming the limitations of the RADseq approach. This can ensure the integration of RADseq data into existing SNP array datasets and enhance their applicability for genomic prediction.

### Performance of phenomic prediction in apple

The concept of phenomic prediction has been proposed as a low-cost alternative to genomic prediction for complex traits and has shown the potential to outperform genomic prediction in terms of predictive ability (Rincent et al., 2018; Zhu et al., 2021; Brault et al., 2022). Our implementation of phenomic prediction models initially focused on identifying the best-performing NIRS pre-processing method. Among the various phenomic prediction scenarios, the one using the first derivative of normalized NIRS had the highest average predictive ability across traits (Fig. S5 in Online Resource 2). This pre-processing method has also demonstrated its superiority in grapevine and has been commonly applied to different crops (Rincent et al., 2018; Zhu et al., 2021; Brault et al., 2022).

The best-performing phenomic prediction scenario showed a strongly reduced average predictive ability across traits compared to both genomic prediction and combined prediction scenarios (Fig. 6). Additionally, the combined prediction did not outperform genomic prediction. At the same time, the best-performing phenomic prediction scenario, which integrated the first derivative of normalized NIRS with the response vector comprising adjusted tree values from the site and year of the NIRS measurement, demonstrated its potential for accurately predicting productivity traits. However, the leaves for the NIRS measurements were collected only after the onset of flowering and fruit development. These biological processes likely altered the chemical composition of the leaves, enabling the phenomic prediction models to accurately distinguish trees based on their productivity of flowers and fruits in the same season. Despite the high predictive ability of these productivity predictions, their applicability is very limited, as they only reflect past biological processes and do not allow for future productivity assessment.

Productivity traits such as flowering intensity in apple are known to be strongly influenced by genotype-by-environment interactions (Jung et al., 2022). To minimize the effect of these interactions in grapevine, Brault et al. (2022) conducted NIRS measurements across different seasons, thereby improving the genetic signal captured by NIRS. Although phenomic prediction models have shown high predictive ability in annual crops (Rincent et al., 2018; Zhu et al., 2021), their application in perennial crops such as apple seems more challenging due to the need to predict phenotypes several years in advance.

Considering the outcomes of our study on genomic prediction of biparental families (Fig. 4), the predictive abilities of phenomic prediction models estimated by LOFO would likely be lower than those reported here using cross-validation. This additional decrease in predictive ability could bring the already low values close to zero, making phenomic selection unsuitable for predicting quantitative apple traits. While the primary interest of this study was to assess the predictive ability for biparental families, the chosen approach to NIRS measurement required substantial resources for the necessary fine leaf milling. This led to the restriction of the NIRS dataset to the apple REFPOP accessions, which did not enable the LOFO validation concept to be tested. Although fine milling allowed for precise NIRS measurements, it diminished the expected cost-effectiveness of phenomic prediction.

### Recommendations for the implementation of genomic selection in apple breeding

Our study demonstrates that the apple REFPOP alone can be used to accurately predict diverse biparental families. Predictive ability for these families can be slightly improved by extending the training set of the apple REFPOP with additional related germplasm. For genomic prediction model training, the size of the apple REFPOP could potentially be reduced by excluding accessions unrelated to the predicted biparental families, thereby saving phenotyping costs—an aspect that remains to be tested.

Predictive ability for the same biparental families can vary depending on factors such as their phenotypic variability, family size, and the validation approach used. Our study recommends applying the validation approach LOFO2 because it compensates for small family size by pooling families and, unlike cross-validation, allows assessment of predictive ability for families with completely unknown phenotypic information, as is the case in practice.

Of the two approaches tested to decrease genotyping costs—RADseq as an alternative to SNP arrays and phenomic prediction based on NIRS as an alternative to genotyping—only RADseq demonstrated sufficiently accurate genomic predictions. Therefore, we recommend its practical application when genotyping additional biparental families. However, this approach must be used in combination with genotype imputation based on high-density SNP array datasets to ensure proper dataset integration and to achieve predictive ability comparable to that of SNP array datasets. Furthermore, the cost of genotyping, particularly library preparation prior to sequencing, needs to be further reduced to improve the applicability of RADseq for genomic selection.

## Supporting information

Supplemental methods - Online Resource 1

Supplemental tables - Online Resource 3

Supplemental figures - Online Resource 2

## STATEMENTS AND DECLARATIONS

### Funding

This study was funded by the Swiss Federal Office for Agriculture project “Apfelzukunft dank Züchtung” (2020/17/AZZ).

### Competing Interests

The authors have no relevant financial or non-financial interests to disclose.

### Author contributions

M.J., A.P., and G.B. conceived the study. M.J., M.H., A.K., and D.K. contributed to data collection. M.J., M.H., D.K., and M.N. performed the data analysis in consultation with S.B.-S., B.S., A.P., and G.B. M.J. wrote the manuscript with support from A.K., M.N., and A.P. All authors provided critical feedback on the manuscript and read and approved the final version for publication.

### Data availability

The phenotypic, genomic, and near-infrared spectroscopy data acquired in this study are available in the ETH Research Collection at [to be updated upon article release]. All previously generated phenotypic and genomic data have been deposited in the INRAe dataset archive at https://doi.org/10.15454/VARJYJ, https://doi.org/10.15454/IOPGYF and https://doi.org/10.15454/1ERHGX.

## Acknowledgements

The authors thank the field technicians at the locations of the participating breeding programs for the maintenance of the orchards and phenotypic data collection. We thank Ingrid Stoffel-Studer, Daniel Ariza Suarez, Steven Yates, and the Genetic Diversity Center at ETH Zurich, Switzerland, for their support with the RADseq laboratory and bioinformatics analysis. We extend our gratitude to Martin Zuber from Agroscope, Switzerland, for assistance with the NIRS measurements. We are grateful to Nick Howard from Fresh Forward, the Netherlands, for providing extensive pedigree information. This study was funded by the FOAG project “Apfelzukunft dank Züchtung” (2020/17/AZZ).

## Notes

### Competing Interest Statement

The authors have declared no competing interest.

