## Supplemental methods - Online Resource 1 for "Evaluation of genomic and phenomic prediction for application in apple breeding"

### Leaf sampling and DNA extraction

#### a. Sample collection

Fresh leaf discs were sampled using a biopsy puncher (6 mm diameter) to obtain four discs, corresponding to approximately 100 mg of dry material. The discs were immediately placed into 1.2 ml 8-Strip Reaction Tubes (E1720-8000, Starlab, Hamburg, Germany) containing three metal beads (2 mm diameter, Uiker, Freienbach, Switzerland) and frozen at -20°C overnight. The following day, the samples were freeze-dried overnight (24 h) using a freeze-drying unit (Alpha 1-2 LD plus, Martin Christ, Osterode am Harz, Germany) to ensure complete dehydration.

#### b. Sample preparation, lysis and binding to the beads

Lyophilized leaf material was ground to a fine powder using TissueLyser II (Qiagen, Hilden, Germany) for 30 seconds twice, shaking at frequence 16.1 Hz. Afterwards, a customized LGC Kit (NAP44500 sbeadex™ test kit * customized ETH Zurich, LGC Genomics GmbH, Berlin, Germany) for DNA extraction was used. An amount of 300 µl of PN lysis buffer, containing 1 µl Proteinase K (PROK-50, Omega Bio-tek, Norcross, USA), was added to each sample tube. The samples were then homogenized by shaking for 5-10 seconds at 16.1 Hz. The homogenized tissue was incubated at 60°C for 60 minutes to facilitate cell lysis. The lysed samples were then centrifuged at 2500 g for 10 minutes to pellet the cellular debris. A 50 µl aliquot of the lysate was transferred to each tube containing 120 µl of Binding Buffer PN and 10 µL of sbeadex™ particle suspension (resuspended before use by vigorous shaking). The samples were then incubated at room temperature for 4 minutes to allow binding to the beads and loaded onto a robotic platform for DNA extraction.

#### c. Automated processing, DNA extraction and quantification

DNA was then extracted using the robotic KingFisher™ Apex & Flex Purification System (Thermo Scientific, Waltham, USA). After the binding process, an initial washing step was performed using 200 µl of Wash Buffer PN1, followed by an additional washing step using 200 µl of Wash Buffer PN1 containing RNAse at a concentration of 4 mg/ml. This was followed by a washing step with 200 µl of Wash Buffer PN2. For the elution, 55 µl of AMP Elution Buffer was used, filled up in a KingFisher 96-well elution plate. After extraction, 22 µl undiluted DNA were mixed to 3 µl of water and 1 µl of gel loading buffer and separated on a 0.6% agarose gel (TAE). Additionally, sample DNA concentration was measured using the Qubit DNA HS assay on a plate reader (Spark 10M Multimode Microplate Reader, Tecan, Maennedorf, Switzerland) in a 384-well black/clear bottom plate. For the assay, 47 µl buffer with 5% dye and 2 µl sample were mixed. Additionally, a standard curve was created using two replicates of 0.5, 10, 25, 50, and 100, with 2 µl of each standard. The excitation wavelength was set to 485 nm and the emission wavelength to 535 nm. The gain of 10 was calculated from the well with the highest concentration. The DNA concentration of the samples was then normalized to a concentration of 5 ng/µl using a pipet robot (PIPETMAX 268, Gilson, Middleton, USA), using milli Q-water as diluent, for a final volume of 10 µl per well. Diluted samples were pipetted into 96-well plates such that each plate included one randomized water control and two positive controls of previously genotyped apple leaf samples.

**ddRAD libraries construction**

**d. Adapter annealing**

EcoRI adapters (Ad Eco-P1.1 and Ad Eco-P1.2) and TaqI adapters (Ad Taq-P2.1 and Ad Taq-P2.2_bio) were prepared separately by mixing 4 µl of each oligonucleotide (100 µM stock) and 2 µl annealing buffer (5x). Annealing buffer is prepared by mixing 50 µl of 10 µM buffer with 50 µl $H_2O$, containing following reagents: 500 mM NaCl, 100 mM Tris/Cl with a pH of 8. The EcoRI adapters were diluted to 0.3 µM using 1x annealing buffer, while TaqI adapters were used directly at 40 µM. Adaptors were then heated to 98°C for 2.5 minutes, followed by cooling down at 2°C per minute until reaching room temperature to allow hybridization.

**e. DNA digestion**

For DNA digestion, the restriction enzymes EcoRI-HF and TaqI-v2 were used. For each 96-well plate, a mastermix was prepared, mixing 550 µl rCutSmart™ Buffer (B6004S, New England Biolabs, Ipswich, USA), 440 µl deionized water, 88 µl EcoRI-HF (R3101S, New England Biolabs, Ipswich, USA), and 88 µl TaqI-v2 (R0149S, New England Biolabs, Ipswich, USA). Then, 5 µl mastermix were added to 10 µl of diluted sample DNA in each well. The thermal cycling conditions were set using a programmable thermal cycler (Labcycler 48s, SensoQuest, Göttingen, Germany) as follows: an initial denaturation step at 37°C for 30 minutes for enzyme activation of EcoR-HF, followed by an incubation at 65°C for 30 minutes to allow TaqI-v2 to digest DNA at their respective recognition sites. Enzyme inactivation was achieved by incubation at 80°C for 20 minutes. Finally, samples were stored at 4°C.

**f. DNA ligation**

For each 96-well plate, a ligation mix was prepared by mixing 330 µl of rATP (10 mM), 220 µl of Taq-P2-biotin Adapter (3 µM), 88 µl of T4 Ligase Buffer (10x, New England Biolabs, Ipswich, USA) and 110 µl of T4 Ligase (400 U/µl, New England Biolabs, Ipswich, USA). Additionally, 1 µl EcoRI P1 adapter (0.3 µM) was added to the digested DNA for each reaction. For the ligation reaction, 6.5 µl of the Ligation Mix was dispensed into each well containing 15 µl of digested DNA. The reaction mixtures were incubated at 23°C for 20 minutes to facilitate ligation, followed by a heat inactivation step at 65°C for 10 minutes.

**g. Sample pooling and size selection**

A total of 48 individually barcoded DNA samples (21 µl each) were pooled in a 2 ml Eppendorf tube and mixed thoroughly. An aliquot of 150 µl from the pooled sample was taken for size selection using AMPure beads (Beckman Coulter, Brea, USA), while the remainder of the pool was stored at -20°C. For the preparation of a 550 bp library, 90 µl of AMPure beads were added to the 150 µl of pooled, ligated DNA (ratio 0.6x), resulting in a total volume of 240 µl. This mixture was pipetted gently to ensure proper mixing and incubated at room temperature for 10 minutes. The tube was then placed on a magnetic stand to separate the beads from the solution, and after 4-5 minutes, the supernatant was carefully saved and transferred to a new tube. Next, 16.10 µl of AMPure beads (0.07x) were added to the 230 µl of supernatant. The mixture was incubated at room temperature for another 10 minutes and then placed on a magnetic stand for separation. After 4-5 minutes, the supernatant was removed and discarded. The beads were washed twice with 300-400 µl of 70% ethanol, ensuring that the ethanol came into contact with the beads each time. The tube was kept on the magnetic stand during the washes. After the final wash, the beads were allowed to dry for 10 minutes. To elute the purified DNA, 30 µl of water were added to the dried beads and mixed. Following a 2-minute incubation at room temperature, the tube was placed on the magnetic stand again, and the purified DNA was transferred to a new tube.

#### h. Selection for P2-biotin labeled adapters

Shortly before use, 15 µl of Dynabeads M-270 Streptavidin (Invitrogen, Waltham, USA) were washed three times with 100 µl of 1xB&W buffer. After washing, the beads were resuspended in 30 µl of 2xB&W buffer. The size-selected DNA (30 µl) was added for DNA binding to the washed bead suspension, resulting in a total volume of 60 µl. This mixture was incubated at room temperature for 15 minutes, with the tube being placed on a thermomixer (Eppendorf ThermoMixer C, Eppendorf, Hamburg, Germany) at 500 rpm and 22°C to ensure vigorous mixing. After the incubation, the tube was placed on a magnetic stand for 2 minutes to separate the beads (bound to the P2-biotin labeled adapters) from the supernatant, which was then discarded. The beads were washed three times with 100 µl of 1xB&W buffer. During the first wash, the beads were mixed with the buffer, while the subsequent two washes were performed with the tube remaining on the magnetic stand to ensure efficient washing and removal of unbound components. Finally, the washed beads were resuspended in 45 µl of distilled water to elute bound DNA from the beads.

#### i. PCR amplification and cleanup

The PCR amplification was carried out using the bead suspension and specific primers. The reaction setup for each of the four tubes included 45 µl of bead suspension, 3 µl of primer 1 (10 µM) from Plate 1 (P195) or Plate 2 (P193), and 3 µl of primer 2 (10 µM) from Plate 1 (P289) or Plate 2 (P285). Additionally, 50 µl of KAPA HiFi HotStart ReadyMix (KK2602, KAPA Biosystems, Wilmington, USA) was added to each tube. The PCR conditions were set as follows: an initial denaturation at 95°C for 2 minutes, followed by 8 cycles of 98°C for 20 seconds, 65°C for 20 seconds, and 72°C for 30 seconds. After the PCR cycles, the reactions were again combined. Since the magnetic beads were included in the PCR reaction, the combined mixture was placed on a magnetic stand to separate the beads. The supernatant of 88 µl was carefully transferred to a new tube for sample cleanup. For this, 57.2 µl of AMPure beads (0.65x) were added to the 88 µl of PCR reaction. The mixture was incubated at room temperature for 10 minutes. For the following incubation, the tube was placed on a magnetic stand for 4 minutes to allow the beads to separate. The supernatant was then removed and discarded. The beads were then washed twice with 300-400 µl of 70% ethanol. The tube remained on the magnetic stand during the washes. After washing, the beads were allowed to air dry for 8 minutes. To elute the clean DNA, 20 µl of water was added to the dried beads and mixed. The mixture was incubated at room temperature for 2 minutes. After this, the tube was placed on the magnetic stand for 2 minutes, and the purified DNA was transferred to a new tube.

#### j. Quality control of the libraries

To assess the quality and size distribution of the DNA fragments, the Qubit fluorometer and the TapeStation (Agilent 4150 TapeStation, Agilent Technologies, Santa Clara, USA) were used following the manufacturers protocol. Molarity was calculated based on the mean fragment size derived from the TapeStation analysis. Specifically, the mean peak height, corresponding to the fragment length from the TS profile, was utilized to convert the DNA concentration from ng/µl (as determined by the Qubit assay) to nanomolar (nM). For subsequent pooling, a volume of 50 µl per pool was required, with two plates being prepared. The Qubit concentration threshold was set at >0.5 ng/µl. For downstream applications, the sample concentrations needed to range between 2 nM and 30 nM, as verified by quantitative PCR (qPCR). This ensured that the samples were within the optimal concentration range for the intended experimental protocols.
