## Supplemental figures - Online Resource 2 for "Evaluation of genomic and phenomic prediction for application in apple breeding"

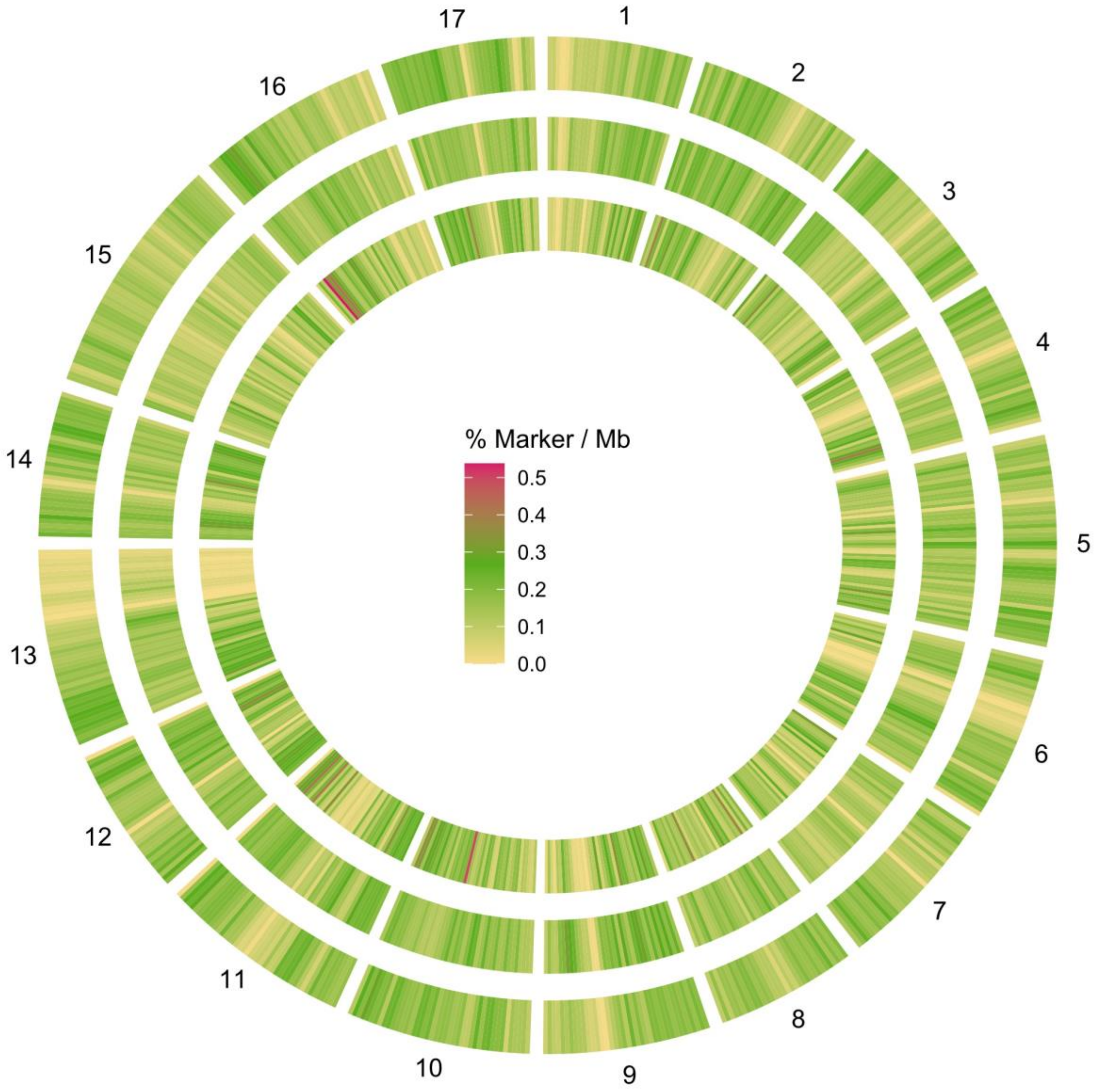


**Fig. S1** Circular heatmap visualizing the percentage of markers per megabase (Mb) of the apple genome, with the SNP array dataset in the outer circle (303,239 SNPs), the RADseq dataset in the middle circle (281,558 SNPs) and their physical overlap in the inner circle (7,255 SNPs).

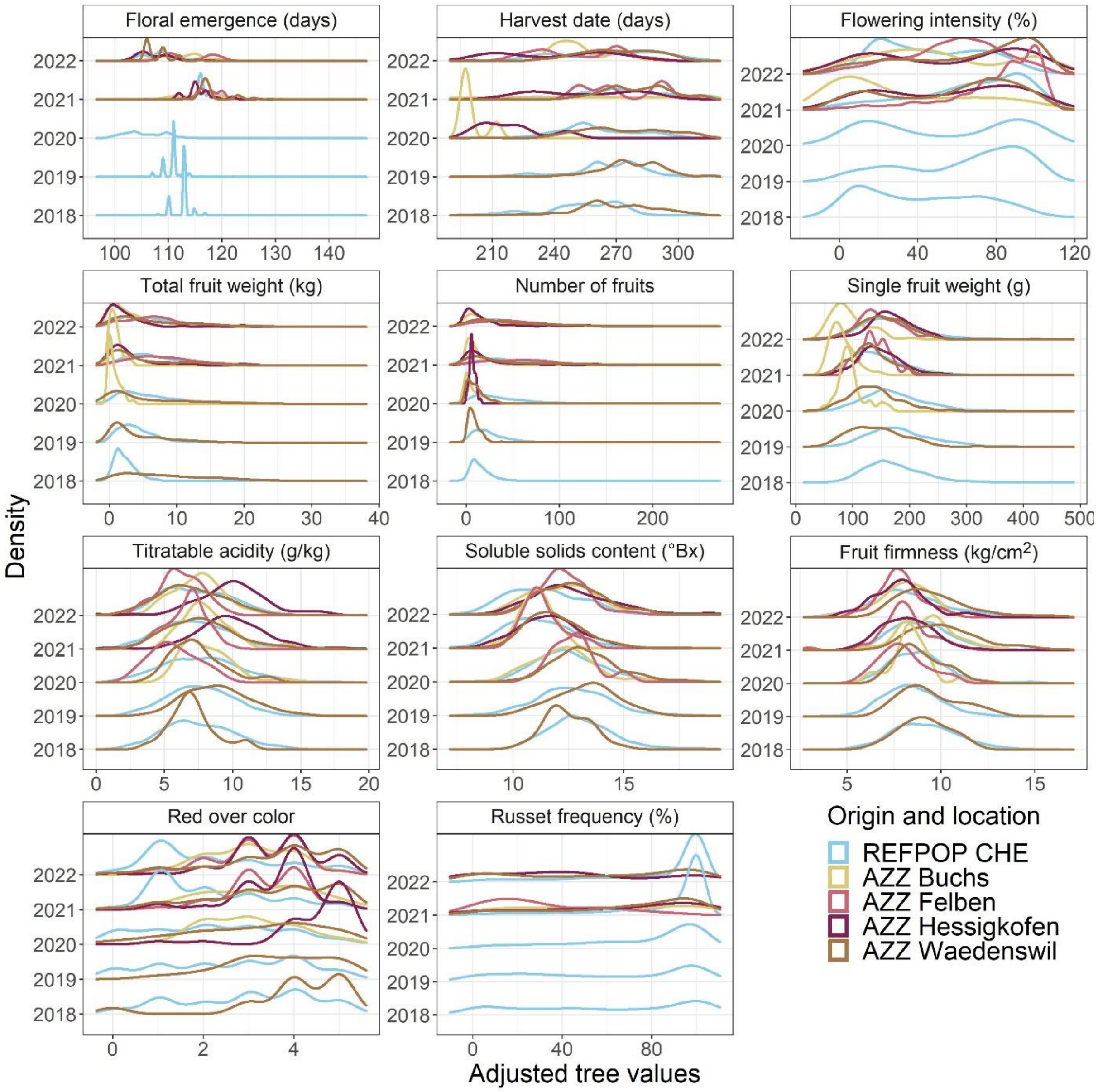


**Fig. S2** Distributions of eleven studied phenotypic traits assessed in Switzerland, shown as density estimates separately for each year of phenotyping, colored by a combination of origin and location of material. The abbreviations stand for: apple REFPOP (REFPOP), AZZ material (AZZ), and Switzerland (CHE).

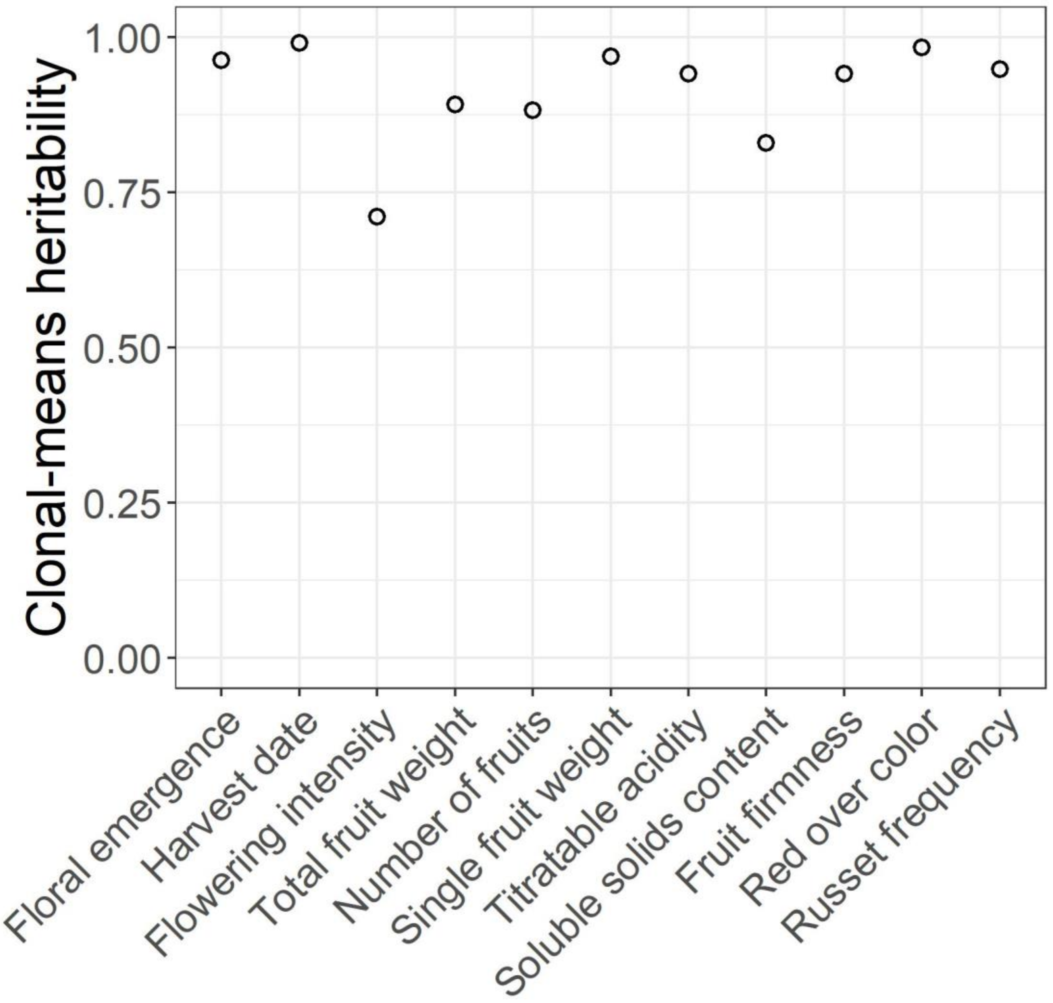


**Fig. S3** Across-environmental clonal mean heritability for eleven traits.

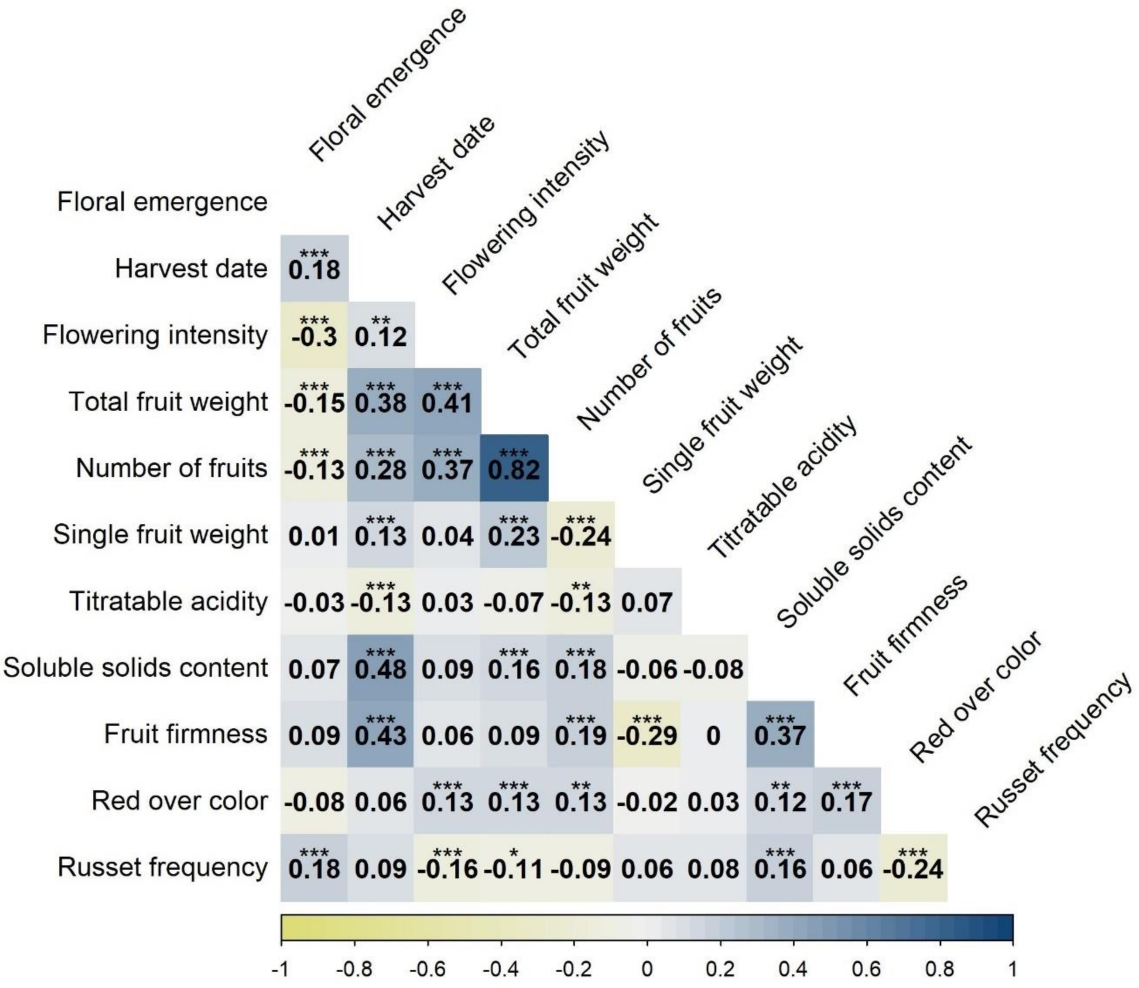


**Fig. S4** Pairwise Pearson's correlations between the clonal values for all pairs of traits. The asterisks indicate significant correlations at Bonferroni-corrected significance levels: * (α = 0.05), ** (α = 0.01), and *** (α = 0.001).

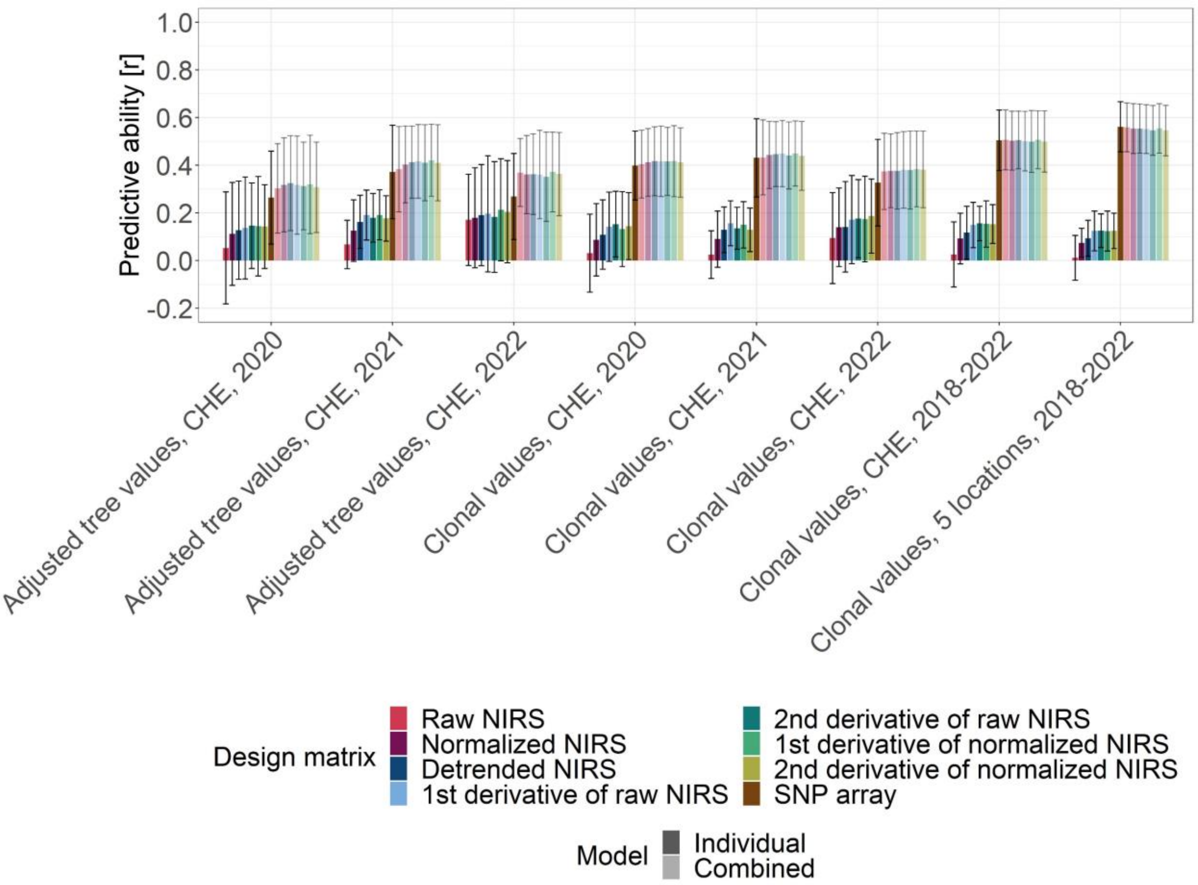


**Fig. S5** Comparison of average predictive ability across eleven traits for phenomic prediction based on near infrared spectroscopy (NIRS) dataset, genomic prediction based on the SNP array dataset, and a combined prediction using both NIRS and SNP array datasets. Predictive ability (y-axis) was assessed by cross validation for different types of design matrices (color) and response values (x-axis, CHE stands for the orchard location in Waedenswil, Switzerland). The individual models implemented random effects using one of the design matrices, the combined models incorporated a combination of the design matrix based on SNP array dataset with one of the design matrices based on NIRS (opacity). Error bars correspond to standard deviation around the mean.
